# USP18 is an essential regulator of muscle cell differentiation and maturation

**DOI:** 10.1101/2022.04.01.486741

**Authors:** Cyriel Sebastiaan Olie, Adán Pinto-Fernández, Andreas Damianou, Iolanda Vendrell, Hailiang Mei, Bianca den Hamer, Erik van der Wal, Jessica C. de Greef, Eleonora Aronica, Vered Raz, Benedikt M. Kessler

## Abstract

The ubiquitin proteasomal system is a critical regulator of muscle physiology and impaired UPS is key in many muscle pathologies. Yet, little is known about the function of deubiquitinating enzymes (DUBs) in the muscle cell context. We performed a genetic screenThe ubiquitin proteasomal system is a critical regulator of muscle physiology and impaired UPS is key in many muscle pathologies. Yet, little is known about the function of deubiquitinating enzymes (DUBs) in the muscle cell context. We performed a genetic screening to identify DUBs as potential regulators of muscle cell differentiation. Surprisingly, we observed that the depletion of ubiquitin-specific protease 18 (USP18) affected the differentiation of muscle cells. USP18 depletion first stimulated differentiation initiation. Later, during differentiation, the absence of USP18 expression abrogated myotube maintenance. USP18 enzymatic function typically attenuates the immune response by removing interferon-stimulated gene 15 (ISG15) from protein substrates. However, in muscle cells, we found that USP18 regulates differentiation independent of ISG15 and the ISG response. Exploring the pattern of RNA expression profiles and protein networks whose levels depend on USP18 expression, we found that differentiation initiation was concomitant with reduced expression of the cell-cycle gene network and altered expression of myogenic transcription (co) factors. We show that USP18 depletion altered the calcium channel gene network, resulting in reduced calcium flux in myotubes. Additionally, we show that reduced expression of sarcomeric proteins in the USP18 proteome was consistent with reduced contractile force in an engineered muscle model. Our results revealed nuclear USP18 as a critical regulator of differentiation initiation and maintenance, independent of its role in the ISG response. to identify DUBs as potential regulators of muscle cell differentiation. Surprisingly, we observed that the depletion of ubiquitin-specific protease 18 (USP18) affected the differentiation of muscle cells. USP18 depletion first stimulated differentiation initiation. Later, during differentiation, the absence of USP18 expression abrogated myotube maintenance. USP18 enzymatic function typically attenuates the immune response by removing interferon-stimulated gene 15 (ISG15) from protein substrates. However, in muscle cells, we found that USP18 regulates differentiation independent of ISG15 and the ISG response. Exploring the pattern of RNA expression profiles and protein networks whose levels depend on USP18 expression, we found that differentiation initiation was concomitant with reduced expression of the cell-cycle gene network and altered expression of myogenic transcription (co) factors. We show that USP18 depletion altered the calcium channel gene network, resulting in reduced calcium flux in myotubes. Additionally, we show that reduced expression of sarcomeric proteins in the USP18 proteome was consistent with reduced contractile force in an engineered muscle model. Our results revealed nuclear USP18 as a critical regulator of differentiation initiation and maintenance, independent of its role in the ISG response.

## Introduction

Muscle degeneration is a hallmark of a wide range of human pathologies that leads to reduced mobility. Muscle degeneration is characterised by atrophy, fat infiltration, fibrosis and impaired regeneration, which are collectively regulated by multiple cellular machineries, among which the ubiquitin proteasome system plays a prominent role ^1^. The muscle-specific E3 ligases Atrogin-1 and MuRF1 play a regulatory role in muscle atrophy by ubiquitinating proteins and thereby tagging them for degradation ^2^. Deubiquitinating peptidases (DUBs) are capable of removing these ubiquitin molecules from substrate proteins and are consequently a regulatory checkpoint in the ubiquitin proteasome system. Sarcopenic muscles were found to be characterised by ubiquitin accumulation, which was concomitant with altered expression and activity of several DUBs ^3^. More recently, USP19 and USP1 were found to be involved in muscle atrophy and possibly also affecting myogenesis ^4,5^. Myogenesis is the formation of new muscle fibres via proliferation and subsequent differentiation of muscle stem cells during muscle regeneration. This process is highly dependent on the timely turnover of muscle-specific regulatory factors ^6^. USP7 and USP10 have been found to regulate myogenesis factors ^7,8^. Together, this indicates that DUBs could contribute to muscle regeneration and degeneration. Despite the large impact of muscle degeneration in pathological conditions, relatively little is known about the role of over 100 DUBs with regards to muscle biology ^9^.

USP18 is a cysteine protease DUB removing interferon stimulated gene 15 (ISG15) from protein substrates (deISGylation) ^10^, hence is a key regulator of the ISG response. Protein modification via ISG15 ^11^, in analogy to ubiquitination, can affect protein stability or localization and thereby alters cellular processes, such as protein translation ^12^, DNA repair ^13^, innate immunity ^14,15^ and antiviral responses ^16,17^. In addition, independent of its enzymatic activity, USP18 can inhibit the type 1 interferon (IFN-1) signal transduction by binding the Interferon-alpha/beta receptor (IFNAR2) ^18^. USP18 expression is induced by several cytokines in immune response, including the type 1 interferons (IFNα and IFNβ). USP18 deficiency causes interferonopathies in mice and in humans ^19–21^. Furthermore, USP18 appears to be involved in tumorigenesis by affecting cell division ^22–24^. A role for USP18 in muscle pathologies is, however, obscure. Unexpectedly, in the hunt for novel regulators of muscle biology, we found USP18 to be essential for muscle differentiation independent of the ISG response.

## Results

### Genetic screen for DUBs affecting muscle cell differentiation

To explore potential novel roles for DUBs during muscle cell differentiation, we transfected immortalised human skeletal muscle cells with a DUB siRNA library. Transfection was allowed to occur in proliferating cells for 48 hours, after which differentiation was induced by serum starvation. After 72 hours of starvation, the differentiated (fused) cells were immuno-labelled with an antibody to Myosin heavy chain (MyHC). Using high throughput imaging the percentage of nuclei in fused cells (fusion index) was quantified (Fig. 1A). In total, siRNA against 40 DUBs led to a reduction in fusion index of more than 2-fold compared with non-transfected differentiated cells (Fig. 1B). Consistent with previous literature ^7,8,25,26^, the knockdown of SENP2, USP7, USP10 and USP13, had a detrimental effect on muscle cell fusion (Fig. 1B-C). Interestingly, knockdown of USP19, which is reported to suppress myogenesis ^27^, did not positively affect the fusion index in this experimental setup (Fig. 1C). siRNA against USP18 was found to reduce the fusion index after 72 hours as compared with control cultures, which has not been reported previously. Knockdown of USP18 and negative effect on MyHC expression was subsequently confirmed by western blot analysis using three different siRNAs (Fig. 1D).

**Figure 1.**
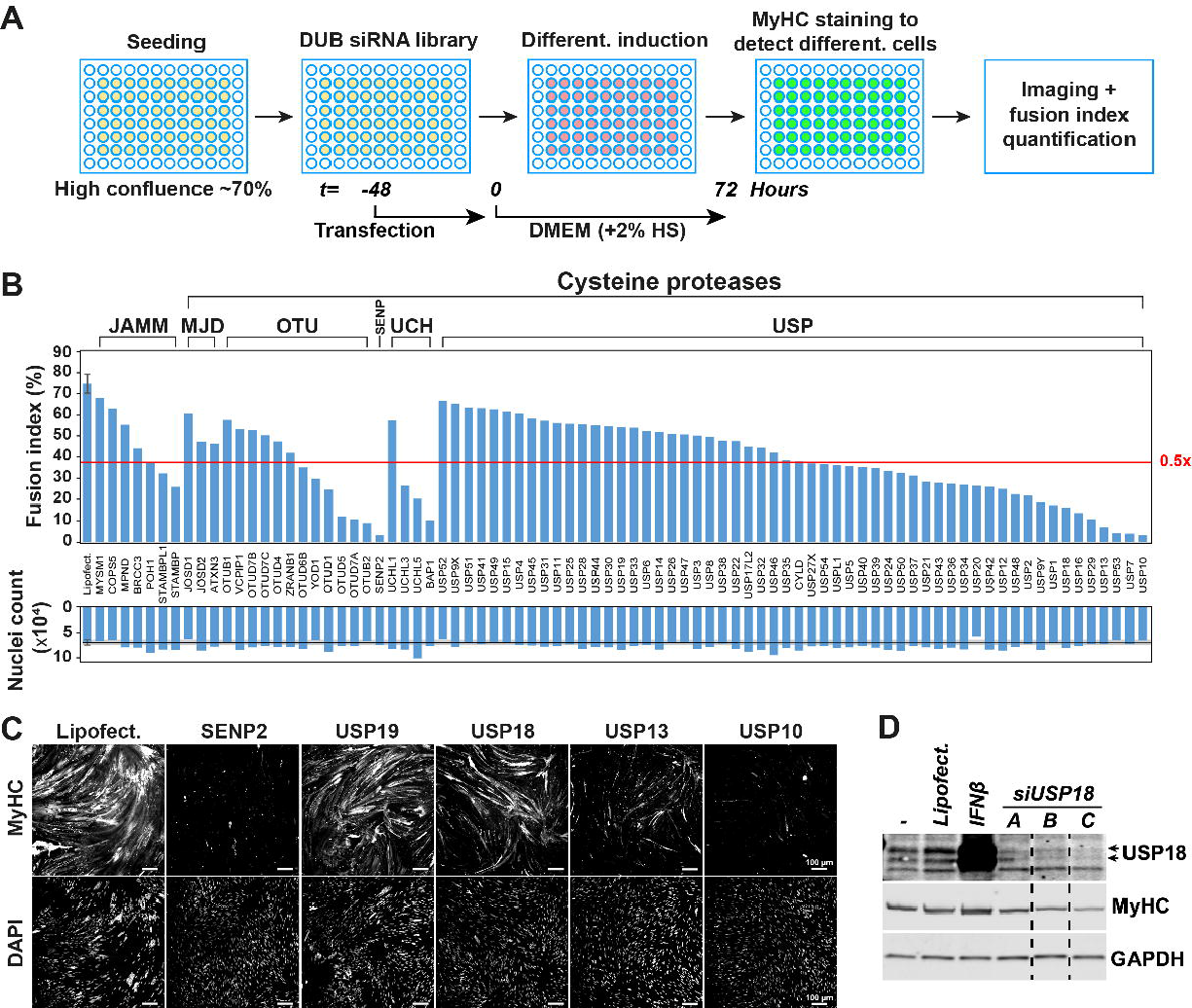
siRNA DUB screen in human muscle cells under differentiation condition. (**A**) A scheme of the screen assay. (**B**) Bar charts show fusion index and nuclei count of the respective DUB siRNA transfections. Red line depicts a 37.5% fusion index cut-off, which is half the fusion index of untreated control cells. (**C**) Representative images show the MyHC and DAPI fluorescent staining for several DUB siRNAs transfections. Scale bar is 100 μm. (**D**) Western blot shows USP18 expression in 72 hours differentiated cell cultures transfected with three different siRNAs to USP18. Cell cultures treated with Lipofectamine or untreated (−) are controls for the knockdown. IFNβ treated cultures were used as positive control for USP18 protein expression. MyHC marks muscle cell differentiation and GAPDH is a loading control.

### USP18 depletion causes accelerated muscle cell differentiation

Standard muscle cell differentiation experiments, including the siRNA screen (Fig. 1A), are carried out in a medium containing 2% horse serum. Horse serum contains exogenous growth factors, hormones and macromolecules, whose exact content remains obscure. Since horse serum could contain factors, like cytokines, that might trigger the IFN-1 pathway and thereby USP18 expression ^28^, we considered excluding horse serum during differentiation. Noticeably, MyHC levels in cultures without horse serum were similar to cultures with horse serum, indicating differentiation (Fig. S1). Therefore, all subsequent differentiation experiments were carried out in serum- and antibiotics- free medium (DMEM −/−, Fig. 2A). During myogenic differentiation, proliferating cells (myoblasts) first commit to differentiation (myocytes) and then fuse and form multinucleated cells (myotubes) that can be recognized by the expression of MyHC (Fig. 2A)^6^. Although USP18 expression is known to be induced by cytokines^28^, noticeably, USP18 levels increased during muscle cell differentiation (Fig. 2B). As USP18 expression was found during the commitment phase, preceding the expression of MyHC (Fig. 2C), we assessed differentiation rates in USP18 KD cell culture at 24 and 72 hours (Fig. 2D). Spontaneous differentiation can be found in highly confluent cultures. Thus, to eliminate spontaneous differentiation at 24 hours, we seeded cells at only 50% confluence before transfection. In contrast to siControl, in USP18 depleted cultures, differentiated cells were found already after 24 hours of starvation (Fig. 2D). After 72 hours, however, the fusion index was significantly reduced in the USP18 KD cell culture (Fig. 2E), consistent with Fig. 1B-C. Knockdown of USP18 was confirmed by immunofluorescence after IFN-1 treatment (Fig. S2A-B).

**Figure 2.**
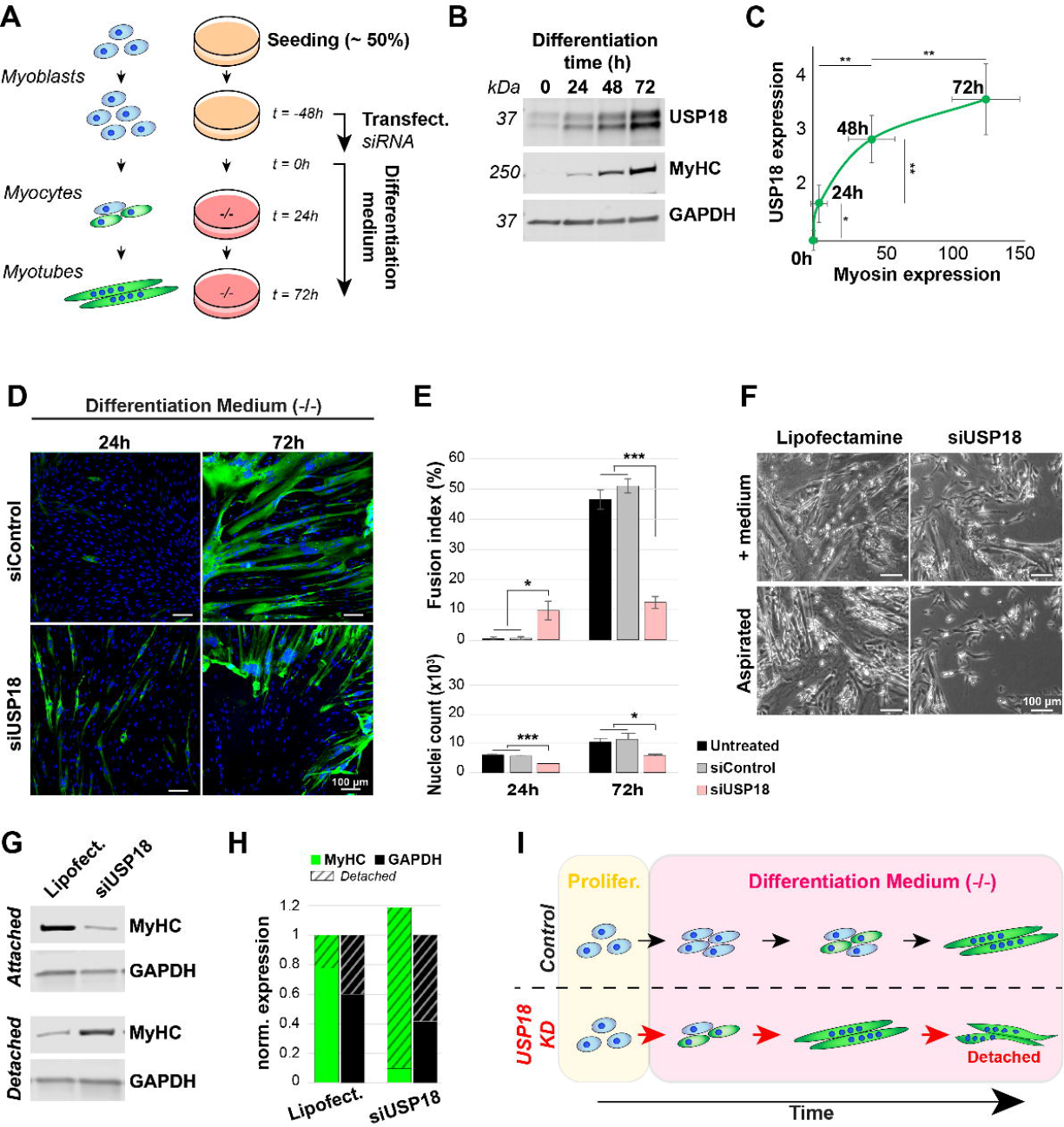
USP18 knockdown (KD) accelerates differentiation in immortalized skeletal muscle cells. (**A**) A schematic overview of muscle cell differentiation and sequential USP18 KD experimental setup. Differentiation in the experiments in this figure were induced by medium starvation without the presence of horse serum, to prevent external factors from triggering USP18 expression. All experiments were performed in at least three biological replicates. (**B**) A representative western blot shows USP18 expression during human muscle cell differentiation marked by MyHC in immortalized cell cultures. GAPDH was used as loading control. (**C**) Scatter plot shows USP18 levels against MyHC levels normalised to GAPDH at 0h, 24h, 48h and 72h of differentiation. Average and error bars (standard deviation (SD)) are from four biological replicates. (**D**) Representative immunofluorescence images show MyHC (green) staining of cell transfected with siControl or siUSP18 cultured in differentiation medium for 24h or 72h. Nuclei are coloured in blue. Scale bar is 100 μm. (**E**) Bar charts show fusion index and cell number of untransfected, siControl or siUSP18 transfected cell cultures of panel D, in respectively black, grey or pink bars. Average and error bars (SD) are from 3 replicates. Statistical significance is depicted by asterisks. (*: 0.05-0.005 and *** <0.0005). (**F**) Images taken with a brightfield microscopy show the differentiated cultures before collection for protein extraction. Scale bar is 100 μm. (**G**) A western blot shows MyHC levels in attached cells after 72h incubation and in the medium collected from the same cultures (=detached) of muscle progenitors after USP18 KD. GAPDH of both fractions is used as loading control. (**H**) Bar chart shows MyHC (green) or GAPDH (black) expression. Attached fraction is in full colour and detached in stripes. MyHC is normalised to the sum of GAPDH in detached and attached fraction. (**I**) Schematic summary of the accelerating effect of USP18 knockdown during starvation (no horse serum) induced differentiation.

Imaging of cell cultures prior to fixation or protein lysis suggested detachment or death of cells (Fig. 2F), which could explain the reduced fusion index after 72 hours of differentiation. To assess this, we generated protein extracts from the still attached cells and the supernatant collected after 72 hours of differentiation. A western blot analysis showed GAPDH and MyHC expression in both fractions (Fig. 2G). As proteins in the supernatant were not degraded, this excludes the possibility of cell death. Quantification showed that ~90% of MyHC levels in USP18 KD cultures was found in the supernatant fraction as compared to only 20% in the supernatant fraction of control cultures. (Fig. 2G-H). This indicates enhanced cell detachment in USP18 KD cells. Together, we suggest that in USP18-depleted cultures, myotubes are formed earlier and detach faster than in control cultures (Fig. 2I).

USP18 is a regulator of cell proliferation that was studied in USP18KD cells ^22,29^. In agreement, a careful cell number quantification showed reduced cell numbers after 24 and 72 hours of differentiation as compared with control cell cultures (Fig. 2E). The increase in fusion index in USP18 KD cell cultures was despite the observed reduced number of nuclei.

To explore whether USP18’s effect on cell differentiation is associated with or independent of cell proliferation, we assessed the effect of USP18 knockdown in the presence of a proliferation medium. We found that proliferation was halted in USP18 KD cell cultures (Fig. 3A) and light microscopy images revealed that myotubes were being formed under proliferating conditions (Fig. 3B). Western blot confirmed that depletion of USP18 triggered MyHC expression in proliferation conditions (Fig. 3C). USP18 KD was confirmed using RT-qPCR, as USP18 protein levels in proliferating cell cultures were under the western blot detection limit (Fig. 3C-D). In addition, USP18 KD in proliferating conditions was also confirmed by immunofluorescence after IFN-1 treatment (Fig. S2C). To substantiate the observation that USP18 KD causes muscle cell differentiation regardless of starvation, we assessed USP18’s effect on differentiation in muscle progenitor cells. Similar to the immortalised muscle cells, USP18 KD induced differentiation after 24 hours under proliferation conditions also in muscle progenitors (Fig. 3E-F and Fig. S3A-B). Noticeably, after 72 hours USP18 KD cultures further differentiated and started to form multinucleated myotubes, whereas the control cells were only beginning to spontaneously express MyHC (Fig. 3E-F and Fig. S4). After 72 hours, cell number in USP18 KD was significantly reduced (Fig. 3F), agreeing with USP18’s role as a positive regulator of cell proliferation. Together, this indicates that USP18 depletion stopped proliferation and induced a commitment switch to myogenic differentiation (Fig. 3G).

**Figure 3.**
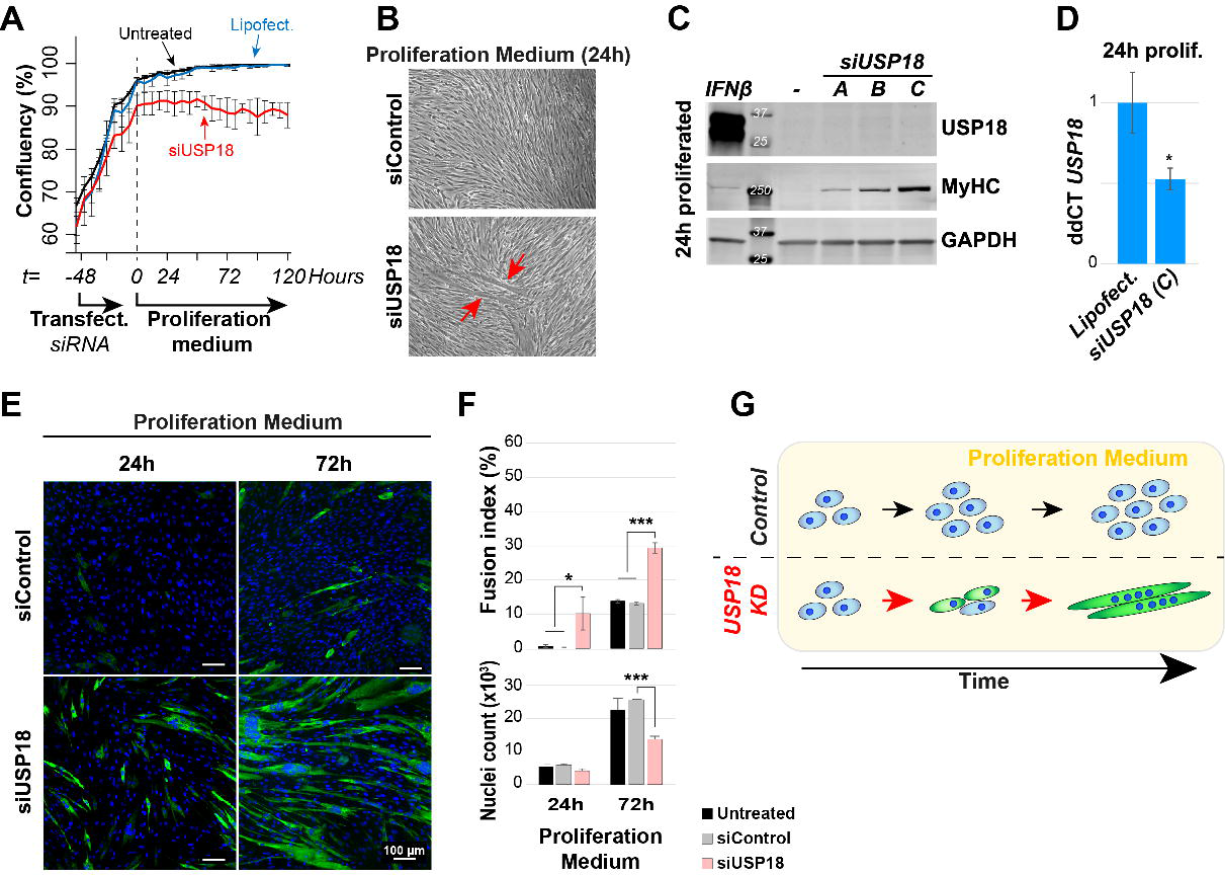
USP18 knockdown (KD) negatively affects proliferation and induces spontaneous starvation-independent differentiation. All experiments in this figure were performed in the proliferation medium. (**A**) Graph shows a change in cellular confluency (%) in USP18 KD cells. Untreated, treated with transfection reagent only or transfected with siUSP18 are depicted with black, blue or red lines, respectively. Cell confluency was measured at 6h intervals. Averages and error bars are from N=3 biological replicates. (**B**) Brightfield microscopy images show transfected confluent cultures of siControl and siUSP18. Red arrows indicate fused cells after 24 hours from medium change. (**C**) Western blot shows increased MyHC expression 24 hours after the transfection medium was replaced with a new proliferation medium with three different siRNAs to USP18. Non-transfected cell cultures (−) is the control for the knockdown. GAPDH is a loading control. In proliferating cell cultures, USP18 is under the detection level. IFNβ treatment was used as positive control for USP18 expression. (**D**) Bar chart shows the average ddCT value of *USP18* gene levels 24 hours after the transfection medium was replaced with a new proliferation medium. Average and error bars (SD) are from N=3. (**E**) Representative immunofluorescence images of control or USP18 KD cells cultured for 24 or 72 hours after the transfection medium was replaced with a new proliferation medium. Myotubes are marked with MyHC (green), and nuclei (blue) mark cell density. Scale bar is 100 μm. (**F**) Bar charts show the fusion index and the nuclear count for the four conditions. Average and error bars (SD) are from N=3 biological replicates. Statistical significance in panel D and F is depicted by asterisks. (*: 0.05-0.005 or *** <0.0005). (**G**) Schematic summary of triggering myogenic differentiation by USP18 depletion regardless of starvation.

### USP18-mediated muscle cell differentiation is independent of deISGylation and IFN-1

As USP18 is known to cleave ISG15 from substrate proteins ^30^, we investigated whether altered deISGylation by USP18 affects myogenic differentiation. We reasoned that an increase in USP18 during muscle cell differentiation might result in increased protein deISGylation. However, in contrast to the elevation in USP18 protein levels, ISG15-conjugates were low in abundance and only showed minor or neglectable changes (Fig. 4A). To further assess whether cell differentiation in USP18-depleted cultures might be dependent on ISG15, we knocked down ISG15 expression using a previously reported siRNA approach ^14,31^. Knockdown in the siISG15 single or siISG15/siUSP18 double transfection was confirmed after IFN-1 treatment (Fig. 4B). ISG15 KD did not affect differentiation, and in the double KD, ISG15 KD did not prevent differentiation driven by USP18 depletion (Fig. 4C-D). This demonstrates that USP18 regulates muscle cell differentiation independent of ISG15. Also, in human muscle progenitor cells, differentiation in the USP18 KD cells was independent of ISG15 and deISGylation (Fig. S5). Together, these data indicate that the USP18 mediated switch between proliferation and differentiation is independent of USP18-dependent deISG15ylation that requires its catalytic activity.

**Figure 4.**
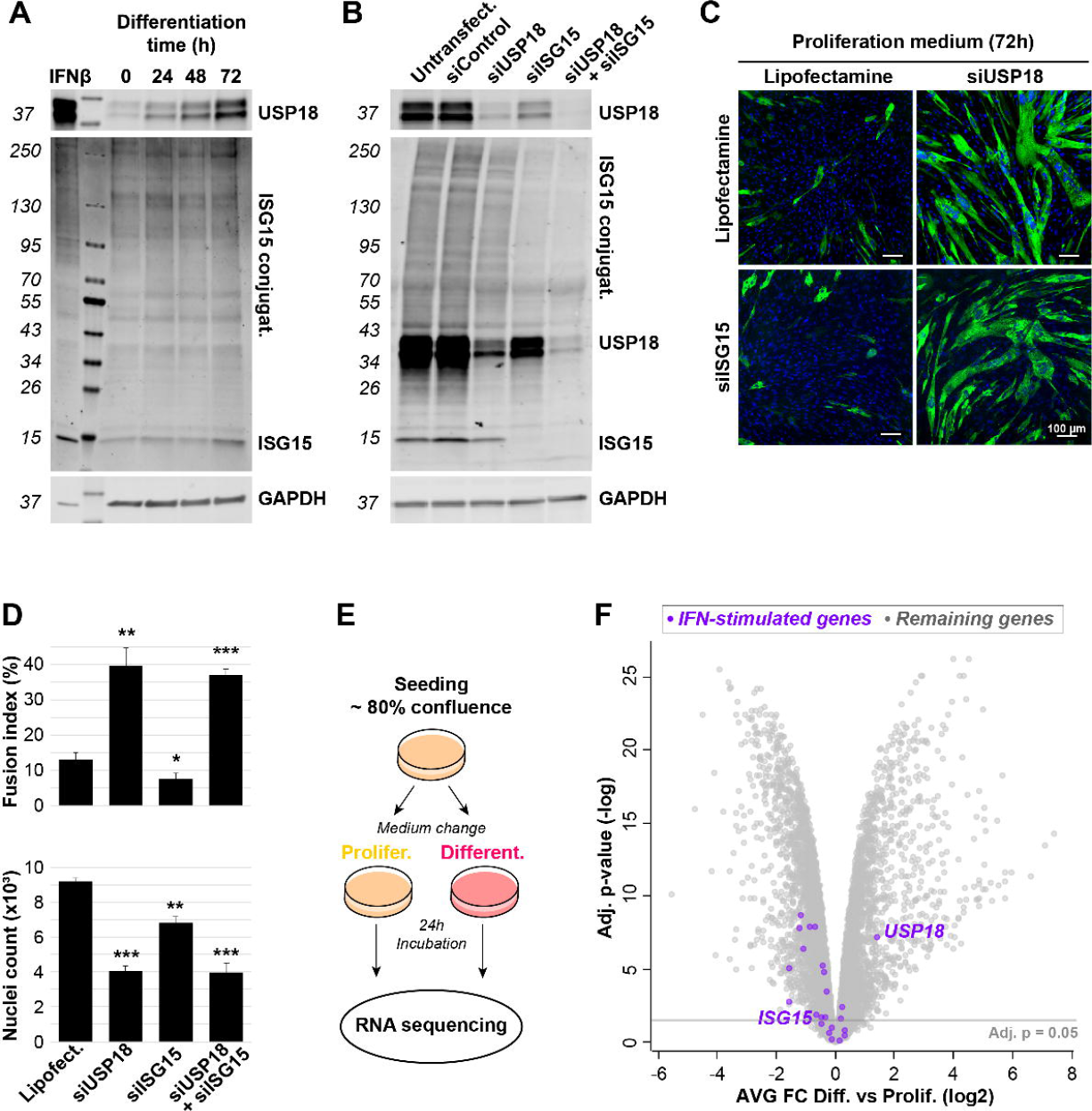
Elevated USP18 expression during muscle cell differentiation is IFN-1-independent. (**A**) Western blot against USP18 and ISG15 during immortalised muscle cell differentiation. GAPDH is used as loading control and IFNβ treatment was used as positive control. (**B**) Western blots show USP18, ISG15 and ISG15 conjugates in untransfected, siControl, siUSP18, siISG15 or siUSP18 + siISG15 double transfected immortalised cell cultures treated with IFNβ treatment (1000U/mL) for 24h in proliferating medium. GAPDH was used as loading control. (**C**) Representative immunofluorescence images show MyHC staining in siUSP18, siISG15 or siUSP18 + siISG15 immortalised cell cultures incubated in proliferation medium for 72h. Nuclei are counter-stained with DAPI (blue). Scale bar is 100 μm. (**D**) Bar charts show fusion index for the respective cell cultures. Average and error bars (SD) are from N=3. Statistical significance is depicted with asterisks. (*: 0.05-0.005, **: 0.005-0.0005, *** <0.0005). (**E**) RNA sequencing experimental setup. Differentiation was induced by medium starvation without the presence of horse serum. (**F**) Volcano plot of transcriptome (N=3) shows mRNA fold change vs. p-value between proliferation and differentiating immortalised muscle cell cultures. The interferon type-1 induced genes are highlighted in purple. The adjusted p-value of 0.05 is depicted.

USP18 expression and other Interferon stimulated genes (ISGs) are typically upregulated by the IFN-1 pathway ^32^. If the IFN-1 pathway is activated during differentiation, we expect to find upregulation of ISGs. However, RNA sequencing of control cell cultures undergoing differentiation showed that only *USP18* is upregulated (Fig. 4E-F). Expression levels of all other ISGs, including *ISG15*, were unchanged or significantly downregulated (Fig. 4F, Table S3). Moreover, both intrinsic *IFN-1-α/β* levels were not detected in the RNA sequencing data, indicating that the ISG response is not activated during differentiation (Table S3). Although USP18 upregulation by other cytokines like TNF-α or type 3 interferons (IFN-III) has been reported ^28^, these cytokines were also not detected in our RNA sequencing data. Together, it is unlikely that these cytokines drive USP18 during muscle cell differentiation, indicating that *USP18* expression may not be driven via canonical cytokine signalling.

### USP18 depletion impacts the myogenic differentiation program

To elucidate myogenic pathways that are initially affected by USP18, mRNA sequencing was performed on USP18-depleted cells that were cultured for 24 hours in a proliferation medium. USP18 depletion led to massive changes in mRNA expression levels (p<0.05, FDR), from which 1445 genes passed the FC>|2| cut off (Fig. 5A). Confirming reduced proliferation and start in differentiation of USP18 KD cell cultures, cell cycle genes were downregulated and myogenic genes were upregulated, respectively (Fig. 5A). To discriminate between the effect of USP18 on differentiation in proliferating *vs*. differentiating conditions, RNA sequencing was performed also on cultures 24 hours after differentiation, and expression patterns that are affected by USP18 were identified by hierarchical clustering on the most affected genes (FC>|4|, N=509). Four prominent clusters were found. The largest cluster (IV, 276 genes) followed the *USP18* expression pattern (Fig. 5B). Cluster IV was significantly enriched with the cell cycle gene network (FDR = 3×10^−68^) (Fig. 5C). Reduced expression of cell cycle genes is consistent with reduced proliferation in USP18 KD cells. The expression pattern in cluster I (80 genes) was opposite to cluster IV and showed increased expression levels in USP18-depleted cultures as compared to control (Fig. 5B). Cluster I was enriched with genes encoding for proteins of the extracellular matrix (ECM), calcium channels and the G-protein coupled receptors (GPCR) (Fig. 5C). Calcium channels and GPRC were also enriched in cluster III (132 genes) showing an expression pattern that is associated with differentiation in both control and USP18 KD (Fig. 5B-C). Calcium channels and GPRC are involved in the myogenic differentiation program ^33,34^. Cluster III also contained muscle contraction gene network, which was also enriched in cluster II (Fig. 5C). However, in cluster II (N=21) their expression was downregulated in USP18 KD cultures compared to control differentiated cultures (Fig. 5B). A comparison of differentiated cultures between USP18 KD and controls showed that the vast majority of the significantly affected muscle structural genes was downregulated in USP18 KD cell culture (Fig. 5D). Together, this suggests that USP18 initially affects differentiation via attenuation of cell proliferation and then affecting the expression of muscle cell differentiation genes.

**Figure 5.**
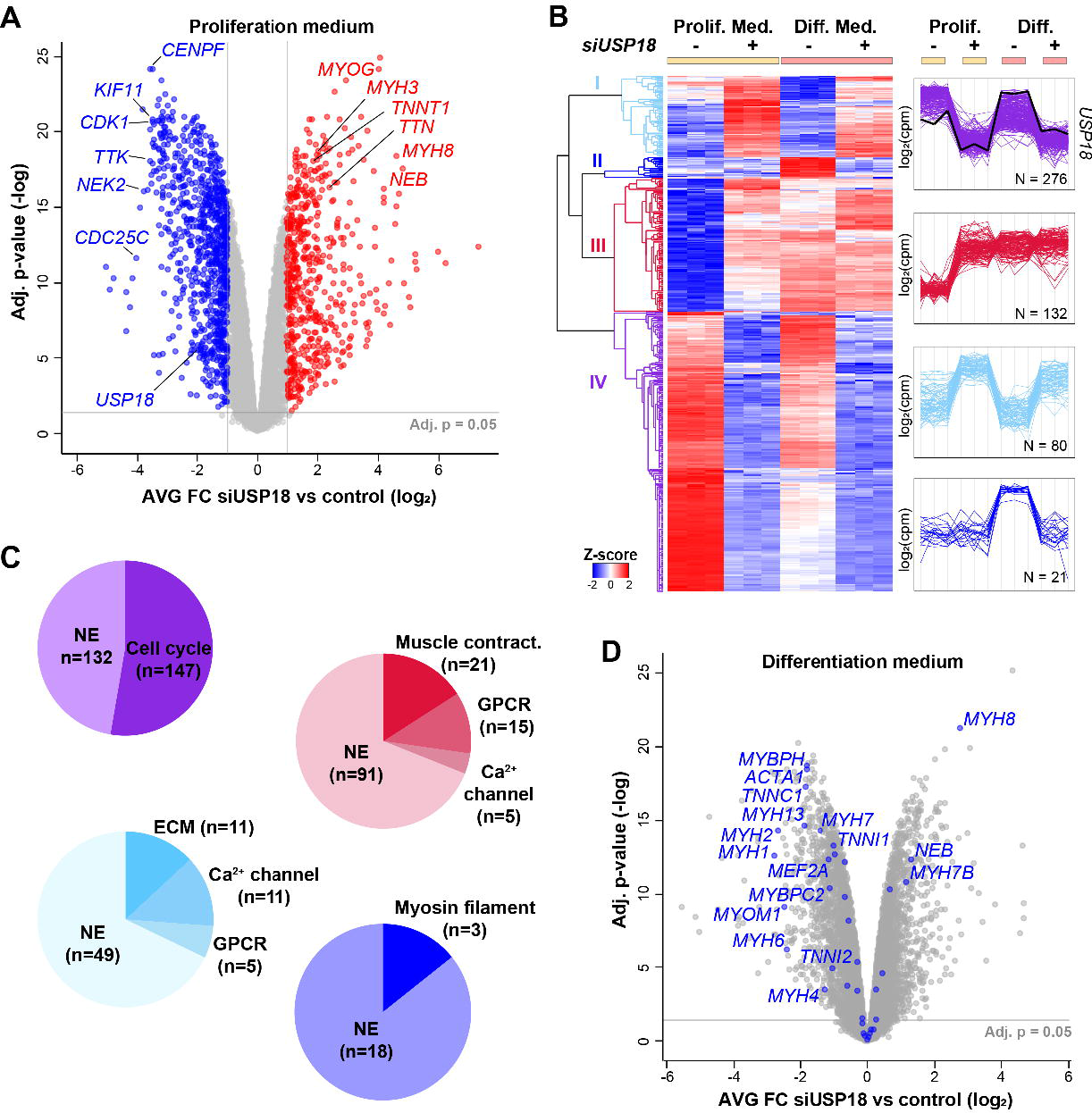
USP18 KD leads to reduced expression of cell cycle genes and dysregulates muscle genes. (**A**) Volcano plot of transcriptome shows mRNA average fold changes in USP18 KD cells versus lipofectamine treated control cells cultured in proliferating medium (N=3). A fold change (FC) cut off (>|2|) and the statistical significance (p<0.05, FDR) are indicated. Genes surpassing these thresholds are depicted in blue (FC<-2) or red (FC> 2). (**B**) Heatmap shows hierarchical clustering of the most affected genes that surpassed the selection criteria (FC>|4|, Adj. p-value<0.05) of the following comparisons: Control vs. siUSP18 cultured in proliferating or differentiating medium. Adjacent plots show expression profiles of the four largest clusters depicted in cyan (I) blue (II), red (III) or purple (IV). The USP18 expression profile is marked with a black line. (**C**) Pie charts show the enriched gene networks per cluster. (**D**) Volcano plot shows mRNA average fold change of USP18 KD cells cultured in differentiation medium for 24h. Myogenic genes are highlighted in blue. Results in this figure were obtained using immortalized cell cultures.

Enrichment in clusters showed a predominant enrichment for cytosolic gene networks. Considering the initial role of myogenic regulatory factors (MRFs) and myocyte enhancing factors (MEFs) during differentiation ^6,35^, we then assessed if gene networks regulating gene expression are affected by USP18 depletion. For this analysis, we first selected genes encoding for nuclear proteins, and then performed enrichment analysis on the 235 genes that were dysregulated (FC>|2|) in USP18 KD cells (Fig. 6A). Enrichment of 36 upregulated muscle-specific transcription regulatory genes was found in USP18 KD cell cultures under proliferating conditions (Fig. 6B). Specifically, *MYOG* levels are initially upregulated during differentiation where *MYF5* and *MYF6* are downregulated ^36^. The expression of these three genes was significantly affected in USP18 KD cell cultures, but showing the same expression pattern as in controls (Fig. 6C). However, the expression pattern of *MYOD1* and *MEF2A* in USP18 KD cells was opposite to control (Fig. 6C and Fig. S6A). RT-qPCR for a subset of these genes validated the RNAseq analysis (Fig. S6A). Protein fold-change of MYOG agreed with the transcript fold change (Fig. S6B). Together, this suggests that differentiation in USP18 KD cells is in part via the canonical myogenic differentiation program, and expression level of these myogenic factors is affected by USP18.

**Figure 6.**
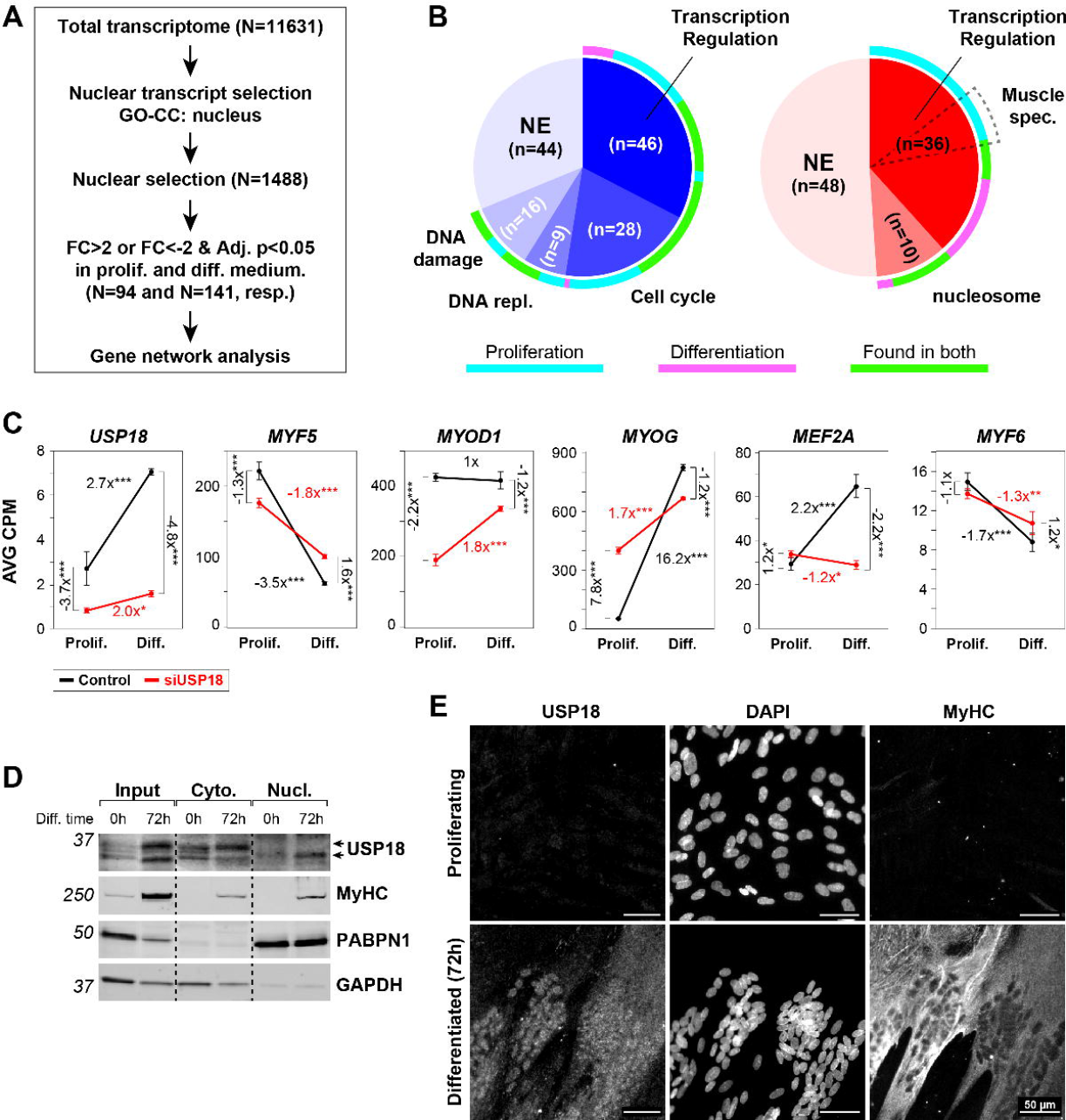
The truncated USP18 isoform is nuclear in differentiated cultures and depletion of USP18 leads to re-programming of myogenic regulatory factors. (**A**) Representative western blot shows cellular localization of USP18, MyHC, PABPN1 and GAPDH in proliferating and 72h differentiated immortalised cell cultures. (**B**) Representative immunofluorescence images show USP18 staining in proliferating and 72h differentiated immortalised cell cultures. MyHC depicts differentiated cells. Scale bar is 50 μm. (**C**) A schematic overview of the nuclear transcript selection. (**D**) Pie charts show significantly enriched UP-keywords in USP18 depleted cultures cultivated for 24 hours. In the left blue pie chart are UP-keywords of genes that are significantly decreased (FC<-2, Adj. p-value<0.05) and in the right red pie chart increased genes (FC>2, Adj. p-value<0.05). The ring around the pies indicate whether the significantly affected genes were found in proliferating (cyan), differentiation (pink) or in both medium conditions (green). Dashed pie depicts the muscle specific genes within the ‘transcription regulation’ UP-keywords. (**E**) Dot plots show the average CPM for five myogenic regulatory genes in control (black) and USP18 KD (red) cells cultured in proliferation or differentiation medium. Average and error bars (SD) are from N=3. Statistical significance is depicted by asterisks. (*: 0.05-0.005, **: 0.005-0.0005, *** <0.0005).

The transcriptome-wide changes caused by USP18 depletion led us to hypothesise that USP18 may be present in the nucleus during cell differentiation. A western blot analysis of subcellular fractions revealed the accumulation of the USP18 34 kDa isoform in the nuclear fraction, whereas the 36 kDa isoform was only found in the cytosol of differentiated cells (Fig. 6D). Immunofluorescence staining in multinucleated myofibers confirmed this observation (Fig. 6E). In agreement, a truncated isoform of USP18 that retains catalytic activity has been found to be expressed in the nucleus ^37^.

If USP18 would affect protein levels, we would expect major changes in the USP18-dependent proteome in differentiated cell cultures. Protein abundance was determined in differentiated cell cultures using label-free liquid chromatography tandem mass spectrometry (LC-MS/MS). Using a high statistical confidence threshold (FDR 0.05), only six proteins were affected by USP18 KD. Considering an FDR of 0.25, altered levels of 63 proteins were significant (Fig. 7A). In total, only 0.18% or 1.88% of the proteins were affected by USP18 in differentiated cells, respectively. This is a minor effect as compared to the USP18-dependent transcriptome. We confirmed differentiation dependent elevation of the skeletal muscle markers MyHC, TNNI1, TNNT3 and KLH31L at the protein level (Fig. 7B).

**Figure 7.**
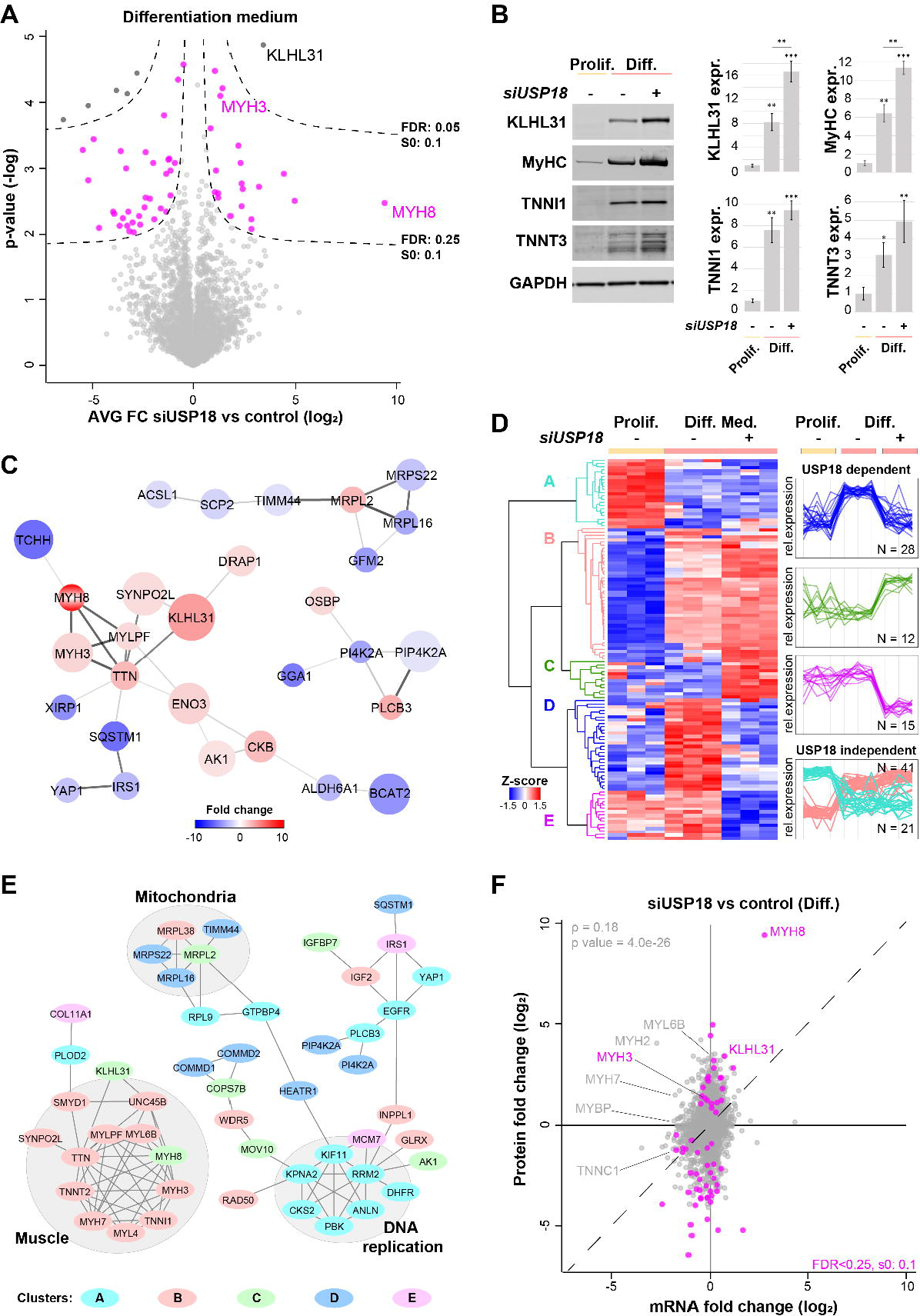
USP18-dependent proteome shows higher expression levels of muscle proteins. (**A**) Volcano plot of USP18-dependent proteome shows the protein average fold changes of USP18 KD cells versus lipofectamine treated control cells cultured in differentiation medium (Diff.) (N=3). Sigmoid curves for FDR 0.05 and 0.25 are depicted by dashed lines. Proteins surpassing the 0.25 FDR threshold are highlighted in pink and in black FDR<0.05. (**B**) Representative western blots show the expression of four muscle-specific proteins. Bar charts show the quantifications for which GAPDH was used as loading control. Expression levels were normalised to control samples in proliferation medium (Prolif.). Average and error bars (SD) are from N=3 biological replicates. Statistical significance is depicted by asterisks. (*: 0.05-0.005, **: 0.005-0.0005, *** <0.0005). (**C**) Protein-protein interaction networks of the proteins that surpassed the FDR<0.25. Colour grade highlights the average fold change, and the larger the circle, the more significant (0.01->0.00001). The protein-protein interaction strength is highlighted by the thickness of the edges. (**D**) Heatmap shows hierarchical clustering for the significantly affected proteins (N=117) of the following comparisons: control vs. siUSP18 cultured in differentiation medium (N=63) and differentiation vs. proliferation medium (N=54). The protein expression profiles of the USP18-dependent clusters are coloured in blue, green or magenta. The cyan and pink clusters represent the USP18-independent changes. (**E**) Protein-protein interaction networks of all 117 proteins, per protein the expression profile cluster is denoted in the corresponding colour. Protein networks are marked by grey circles. (**F**) Scatterplot showing the mRNA against protein fold-change of USP18 KD cells versus lipofectamine treated control cells cultured in differentiation medium. Proteins highlighted in pink pass the FDR<0.25, in grey are the non-significant proteins. Muscle proteins are highlighted. The dashed line indicates the diagonal, and the Spearman correlation is depicted in light grey. Results in this figure are obtained from immortalised cell cultures.

Protein-protein interaction network analysis revealed a USP18 KD-dependent increase in the muscle-specific cluster (Fig. 7C). To identify expression trends that are associated with USP18 KD-driven differentiation, hierarchical clustering was performed on all significantly affected proteins (N=117, Table S4). Five distinctive clusters were found of which the largest cluster (B) consisted of 41 proteins that increased in starved control and starved USP18 depleted cultures (Fig. 7D). 11 proteins were annotated to the muscle network (Fig. 7E), of which nine proteins were found back in the RNA cluster III (Fig. 5B). The opposite expression profiles, decreased expression in both starved control and USP18-depleted cultures, were found in cluster A (N=21). Seven proteins from that cluster were found to be part of the DNA replication network (Fig. 7E), of which six were overlapping with RNA cluster IV (Fig. 5B-C). Cluster D consisted of 28 proteins, including mitochondrial proteins and endosomal proteins that were increased in differentiated control cultures but unchanged in USP18 KD differentiated cultures (Fig. 7D). The expression profiles of cluster C and E showed a steep increase or decrease in USP18 KD differentiated cells as compared to controls (Fig. 7D). Protein-protein interaction network analysis on the 117 significantly affected proteins showed that the muscle proteins, DNA replication, and the mitochondria were the most affected networks (Fig. 7E). For the 63 affected proteins, we compared fold changes between RNA and protein and found a higher change on protein levels (Fig. 7F). This suggests that protein stability might also affected by USP18.

### *In vitro* muscle maturation is disrupted in USP18 depleted cells

Since the sarcomere network and calcium channels were amongst the affected networks in USP18 KD, we assessed whether the function of both networks is impacted. A calcium flux was measured in multinucleated cells. In control cultures, multinucleated cells showed a strong calcium flux, whereas this was absent in undifferentiated cells. (Fig. S7). Calcium flux in cell cultures transfected with siControl was indifferent from parental cell culture counterparts (Fig. 8 and Fig. S7). USP18 depleted multinucleated cells, cultured in proliferation medium, showed a strong induction of the calcium flux (Fig. 8A-B). In contrast, a calcium flux induction was not detected in USP18 depleted multinucleated cells that were formed in differentiating medium (Fig. 8A-B).

**Figure 8.**
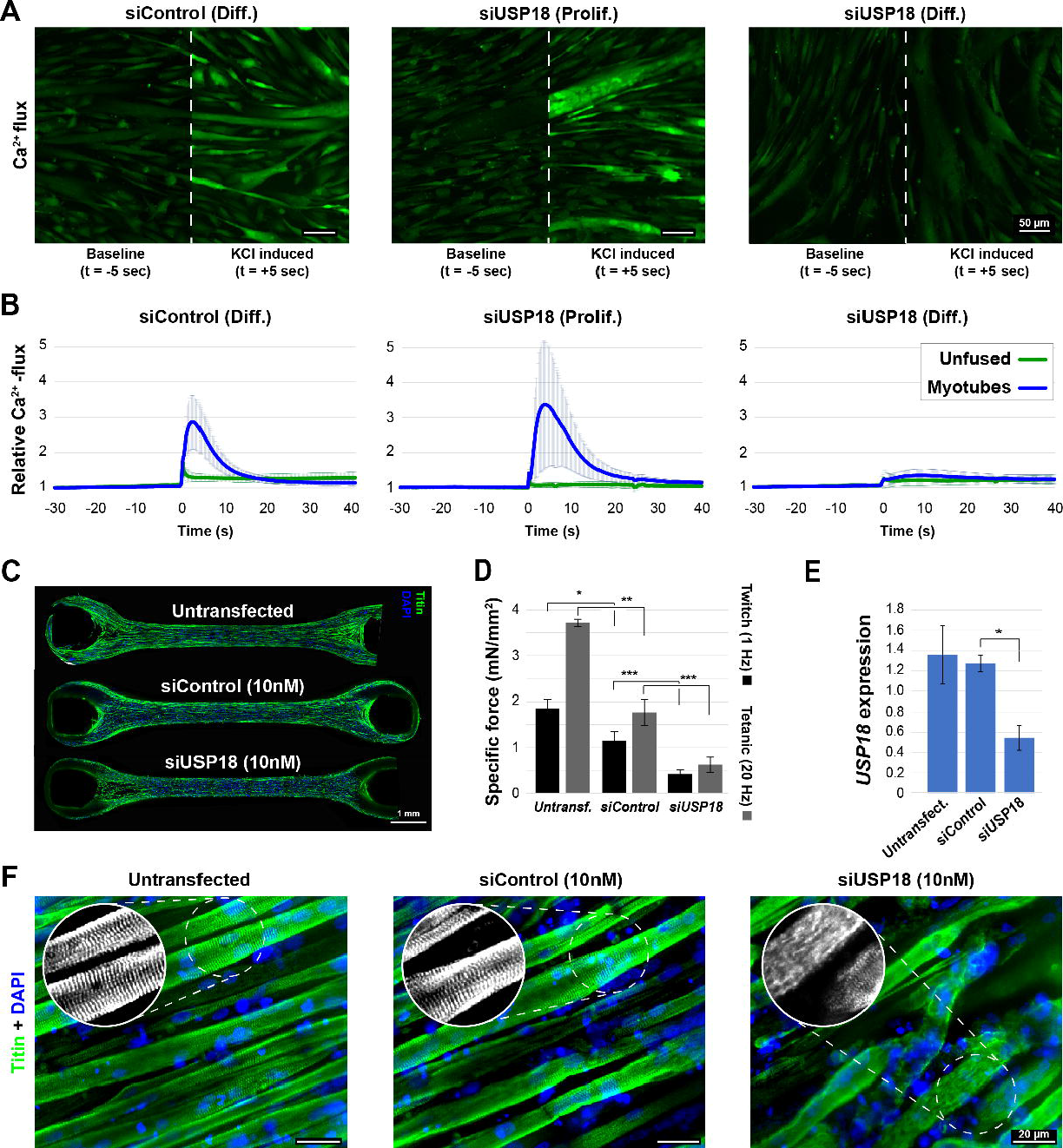
USP18 KD negatively affects *in vitro* muscle physiology. (**A**) Representative images of the calcium flux (green signal) 5 seconds before or after KCl (10 mM) administration in immortalised muscle cells transfected with siControl in differentiation medium (Diff.) or with siUSP18 in proliferation or differentiation medium. The respective images are separated by a white dashed line. Scale bar is 50 μm. (**B**) Graphs show changes in calcium flux over time. Calcium flux in uni-nucleated cells (unfused) is depicted by a green line and multinucleated cells (myotubes) by a blue line. Average and error bars (SD) are from N=3 biological replicates. (**C**) An image of a representative bundle made with muscle progenitor cell line stained for titin (green) and nuclei (blue) for each genetic condition. Scale bar is 1 mm. (**D**) Bar chart shows the specific contractile forces (in mN/mm^2^) at 7 days of differentiation for USP18 KD and controls. Twitch (1 Hz) and tetanic (20 Hz) contractions are depicted by black or grey bars, respectively. Average and SD are from contractions of four different muscle bundles. (**E**) Bar chart shows *USP18* expression levels normalised to untreated muscle bundles at 7 days of differentiation. Averages and SD are from N=3. Statistical significance in panel D and E is depicted by asterisks. (*: 0.05-0.005, **: 0.005-0.0005, *** <0.0005). (**F**) High resolution images show titin architecture in muscle bundles after 7 days of differentiation. The coloured images show titin in green and nuclei in blue. Grey scale images show titin staining only. Scale bar is 20 μm.

We then investigated the effect of USP18 on muscle contraction in 3D muscle bundles. Intact muscle bundles were formed after siRNA transfection (siControl or siUSP18, Fig. 8C). In USP18 depleted muscle bundles, both twitch and tetanic contractile forces were reduced as compared to control siRNA transfected muscle bundles (Fig. 8D). A USP18 KD in the muscle bundle was confirmed using RT-qPCR (Fig. 8E). The structural integrity of the muscle bundles was assessed by staining for titin. In USP18 depleted muscle bundles, a clear disruption in myofibers continuity was found as compared to control muscle bundles, which explains the reduced contractile force (Fig. 8F). Moreover, a very weak striated titin staining was found in USP18-depleted myofibers as compared to controls, indicating sarcomeric disruption (Fig. 8F). This suggests that muscle fibres are formed but not maintained, which is consistent with USP18 KD-mediated detachment of fully differentiated cells in 2D cultures using differentiation medium without serum (Fig. 2D-H).

## Discussion

USP18 has been studied predominantly in the context of immune responses. In this scenario, USP18 deISGylation activity plays a regulatory role in the ISG response ^14,32,38,39^. Here, we reveal that during muscle cell differentiation USP18 plays a role independent of the ISG response. We found that USP18 expression is upregulated during differentiation, but the expression of other ISG genes was unchanged or downregulated. During differentiation or in USP18 depleted cells, ISGylation was not affected. Furthermore, we also show that ISG15 KD had no impact on differentiation nor did it prevent USP18 KD-mediated differentiation. In agreement, in mouse C2C12 cells, ISG15 KD did not affect muscle cell differentiation ^40^. Together, this suggests a non-canonical function of USP18 in muscle cell differentiation.

ISGylation has multiple effects on substrate proteins including protein turnover ^41^. So far, most studies in USP18 depleted cells were focused on global changes in protein levels or ISGylated proteins ^14,42^. During muscle cell differentiation, USP18 depletion had only a small effect on the proteome. Instead, we found a genome-wide impact on mRNA expression profiles in USP18 KD cell cultures. The significant change in transcripts encoding for nuclear proteins, including key myogenic (co)transcription factors in USP18 KD cell culture, agrees with the differentiation phenotype under these conditions.

Similarly, USP18 is a regulator or cell cycle, mediated by its deISGylation activity ^22,29,43,44^. Also, in muscle cells, USP18 depletion leads to reduced proliferation. Yet, it was not associated with global changes in ISGylation, but instead the expression of cell cycle genes was massively downregulated. Based on our results, we suggest that USP18 may act as a safeguard blocking the entry to differentiation, possibly inhibiting the expression of myogenic (co)transcription factors. From our studies we cannot conclude that USP18 directly (co)regulate transcription, however, we found nuclear USP18 in differentiating cells. Nuclear USP18 is associated with ISGF3 transcription factor that bind to ISRE elements ^*45,46*^. Yet, a direct role for USP18 in transcription regulation has not been reported, but it might form a complex with STAT2/STAT1/IRF9 to enhance expression of ISGs in cancer cells ^47^. In analogy to this scenario, we found nuclear accumulation of the truncated isoform of USP18 in differentiating muscle cells, suggesting that this may be linked to triggering differentiation.

Muscle cell differentiation is a multi-step program, where myogenic (co-)transcription factor levels regulate the expression of muscle structural proteins ^6^. Interestingly, we found that USP18 depletion first enhances differentiation, but was concomitant with dysregulation of myogenic transcription factors. Subsequently, at a later stage, the differentiation index is reduced due to cell detachment. The integrity of differentiated myotubes is maintained by sarcomere and cytoskeletal proteins facilitating contraction ^48,49^. Reduced expression of muscle structural (sarcomere) genes was found in USP18 depleted differentiated cells, and accordingly reduced contraction was found in a muscle bundle with USP18-depletion. Specifically, we showed that sarcomere integrity was impacted by reduced USP18 levels. Among the sarcomere genes, *MYOM1* levels were strongly reduced in USP18 KD cultures. MYOM1 is essential for sarcomere stability and crucial for contraction ^50^. In MYOM1 KD cardiomyocytes calcium homeostasis was impaired and contraction was reduced ^51^. Together, our results suggest that USP18 initially blocks entry into differentiation and subsequently regulates the expression of proteins that form and maintain mature myofibers.

Under pathological conditions, USP18 expression is upregulated by interferons, IFN-1 and type III IFN, lipopolysaccharide (LPS) or tumour necrosis factor alpha (TNF-α) ^28^. USP18 expression was also found in Dermatomyositis, a muscular inflammatory disease ^52,53^. In dermatomyositis, both USP18 and ISG15 are upregulated in inflamed muscle tissue regions. In these areas, ISG15 expression/conjugation was higher than in control muscles ^52^. Our results, however, suggest that USP18 plays a separate role in regeneration. In acute injured skeletal muscles, inflammation precedes removal of damaged tissue, which subsequently triggers regeneration via the secretion of cytokines, such as IFN-γ and TNF-α ^54–56^. It is unknown whether IFN-1 signalling is part of muscular inflammation that triggers muscle regeneration. Excessive IFN-1 inflammation could have a causal relation with degeneration/regeneration in dermatomyositis. Engineered muscle bundles combined with inflammatory cells ^57^, could present a relevant model to explore the role of USP18 in myositis. Such a model will provide the framework for therapeutics development. Taken together, our study shows that USP18 function in cell biology may reach beyond the ISG response. We show that USP18 regulates muscle cell differentiation independent of (de)ISGylation. USP18 depletion causes muscle cells to switch from proliferation to differentiation irrespective of serum starvation. USP18 also plays a role in differentiation and maturation, ensuring proper expression of sarcomeric genes. Future research will have to pinpoint the exact molecular mechanism by which USP18 is induced and how USP18 subsequently regulates nuclear processes, such as inducing transcriptional regulators of muscle cell differentiation.

### Limitations of the study

This study describes a novel function for USP18 in myogenesis that appears to be independent of its role in protein deISGylation. This was demonstrated by knockdown studies (USP18 and ISG15 and both together) as well as transcriptomics and proteomics to profile cellular responses including the ISG signature. Removal of USP18 stopped proliferation in human muscle cells and caused a switch to accelerated differentiation, thereby precluding efforts to generate a USP18 knockout or introduce a USP18 catalytically dead mutant. A selective USP18 inhibitor, when becoming available, might represent a route into discriminating between canonical (catalytic) and noncanonical functions, thereby enabling a therapeutic strategy to accelerate muscle regeneration.

## Methods

### Cell culture and siRNA transfection

Immortalized human skeletal muscle cells (7304.1, ^58^) were cultured in growth medium (F10 (Gibco) medium supplemented with 15% FCS, 1 ng/ml bFGF, 10 ng/ml EGF and 0.4 μg/ml Dexamethasone). Differentiation was induced using differentiation medium (DMEM + 2% horse serum or DMEM −/−) for 24h to 72h as specified. Differentiation rate in DMEM −/− medium was assessed by a western blot, and seems to exacerbate detachment on myotubes.

Muscle induced pluripotent stem cells (muscle progenitors ^59^) were cultured in DMEM containing 10% FCS, 1% penicillin–streptomycin and 100 ng/mL bFGF (freshly added). Muscle progenitors were differentiated using DMEM containing 1% Insulin-Transferrin-Selenium-Ethanolamine (ITS-X) and 1% Knockout Serum Replacement (KOSR).

The human ON-TARGETplus siRNA Library (Deubiquitinating Enzymes) (Dharmacon) was prepared in 96-well plate using Lipofectamine RNAiMAX transfection reagent, according to the manufacturer’s protocol (Thermo Fisher Scientific). Cell were transfected at high confluency (~70%) for 48h in transfection medium (80% proliferation medium and 20% OptiMEM) allowing cell density to reach a 90-100% before knockdowns were achieved. Final concentration of siRNA was 10 nM. Differentiation was subsequently induced by switching the medium to differentiation medium with 2% horse serum (Gibco).

In subsequent experiments, siRNAs specific to USP18, ISG15 (Table S1) were transfected into cells according to the manufacturer’s protocol. SiGenome (Dharmacon) was used as control siRNA (named as siControl) (Table S1). Unless specified in the experiments, USP18 KD was carried out with siRNA-C. Cell cultures, at roughly 50% confluence, were transfected with 10 nM siRNA and cultured for 48h in transfection medium (80% proliferation medium and 20% OptiMEM). Subsequently, medium was replaced either with normal proliferation or differentiation medium (DMEM −/−, unless specifically indicated as in the siRNA library screen), or cells were directly harvested according to the experimental setup. Treatments with IFN were carried out with the human interferon IFN-α2 (PBL Assay Science, Cat. No. 11105-1) or recombinant Human IFN-β (Peprotech, Cat. No. 300-02BC) and were used at 1000 U/mL for 24h (unless indicated otherwise). Cell growth rate was determined with the phase contrast channel of the IncuCyte Zoom Imager (Essen Bioscience). An interval of 6h was used to determine the effect of USP18 knockdown on cell growth indicated by the percent confluence over time. All cell culture experiments were performed in the immortalized cell line unless stated otherwise.

### Muscle bundle assay

Muscle bundles were generated as described previously with small adjustments ^49,60^. Muscle progenitor cells were used to create the bundles. Muscle progenitors were trypsinized, after which proliferation medium was added. Cells were subsequently centrifuged and trypsin-containing medium was aspirated. Next, muscle progenitors were diluted in cold growth medium without antibiotics. An ice-cold hydrogel, consisting of 20% Growth Factor Reduced Matrigel (Corning) and 2 mg/ml bovine fibrinogen (Sigma-Aldrich) was combined with muscle progenitors (6×10^5^ cells per bundle). Knockdowns were induced by directly adding the USP18 (or siControl) siRNA-lipofectamine particles at 200 nM to the muscle progenitor cells before adding this to the ice-cold hydrogel mix. Bovine Thrombin (0.8 units) (Sigma-Aldrich) was added to the hydrogel mix before casting the bundle in the polydimethylsiloxane (PDMS) chambers. After full polymerization (after casting 30◻min at 37◻°C), the newly formed bundles were kept in standard 2D muscle progenitor proliferation medium containing 1.5 mg/mL 6-aminocaproic acid (6-ACA) (Sigma-Aldrich). After 48 hours, proliferation medium was changed to differentiation medium (containing 2 mg/mL 6-ACA) and bundles were incubated on a platform shaker at 37◻°C, 5% CO_2_. Half of the differentiation medium was changed every other day. Twitch (1 Hz) and tetanic (20 Hz) contractions were measured after seven days of differentiation using an in-house setup. In brief, a cathode and anode were placed on either one of the PDMS pillars. An electrical pulse at 1 or 20 Hz was applied causing the muscle bundles to contract. Contractile force was calculated based on the displacement of the pillars of one contraction, which was recorded using live microscopy. The specific force was then calculated and normalised to cross sectional area of the muscle bundle. To explore reasons for possible changes in contractile force, bundles were subsequently either fixed overnight in 2% PFA for immunofluorescent staining and stored in PBS at 4◻°C or frozen in liquid nitrogen and stored in −80◻°C for RNA extraction.

#### Protein analyses

##### Western blot analyses

Proteins extracts were obtained using RIPA lysis buffer (20 mM Tris, pH 7.4, 150 mM NaCl, 5 mM EDTA, 1% NP40, 0.1% SDS and 1 mM DTT and 1x protease inhibitor cocktail). Next, 25 μg was separated on SDS-PAGE (Criterion XT, Bio-Rad). Proteins were then transferred to PVDF membranes, which after one-hour blocking (5% milk in PBST) were blotted for the primary antibodies and subsequently with secondary antibodies (Table S2). An Odyssey CLx Infrared imaging system (LiCOR, NE. USA) was used to detect the fluorescent signal. Quantification of protein abundance was done using Image Studio Software version 5.2. Values were corrected for background and normalised to loading controls.

##### Mass spectrometry and analysis

Immortalized muscle cell cultures were harvested by directly adding lysis buffer (20 mM HEPES (pH 8.0), 150 mM NaCl, 0.2% NP40, 10% glycerol and freshly added protease inhibitor cocktail (Roche)) to the cell culture dishes. 100 μg protein lysate was taken for in-solution digestion. First, samples were reduced using DTT (5 mM final concentration) for 30◻min at 37◻°C. Next, iodoacetamide was used to alkylate the samples (final concentration 20 mM, 15 min RT in the dark). Methanol/chloroform extraction method was then used to precipitate proteins, after which proteins were resuspended in 50◻μL of 6◻M urea and subsequently diluted to reduce urea concentration to <1M. Proteins were digested with trypsin (final concentration 0.2 μg/μL) on an overhead rotor at 12 rpm overnight. Digested peptides were subsequently acidified using trifluoroacetic acid (final concentration 1%) and desalted using C18 solid-phase extraction cartridges following the manufacturer’s instructions (SEP-PAK plus, Waters). Reconstituted peptides (in 98% MilliQ, 2% CH_3_CN and 0.1 %TFA) were analysed by nano-liquid chromatography tandem mass spectrometry (LC-MS/MS) using the Ultimate 3000 UHPLC (Thermo Fisher Scientific) connected to an Orbitrap Fusion Lumos Tribrid mass spectrometer (Thermo Fisher Scientific) using a one-hour gradient and universal settings for data acquisition as detailed previously ^61^. Maxquant software (v1.6.10.43) was then used to search the raw MS files against the UniProtKB human sequence database (UP000005640, 79,038 human sequence entries). Next, the following parameters were set to perform label free quantification: Modifications were set on Carbamidomethyl (C) as fixed and Oxidation (M) and Deamidation (NQ) as variable and a maximum of two missed cleavages were allowed. Match between runs function was not used. The quantified proteome was then analysed using Perseus (v.1.6.14.0), where significance was assessed using a student’s t-test with multiple testing using permutation false discovery rate (FDR) set on 0.05 or 0.25 and s0 = 0.1, since the cultures were strongly heterogenic (less than half was fused).

##### Immunofluorescence staining and image quantification

Cell cultures were fixed (10 min, 4% PFA), permeabilized (10 min at −20◻°C with ice cold MeOH), blocked (1h, 5% BSA) and stained using antibodies against USP18, ISG15 and MyHC (Table S2). USP18 antibody was incubated overnight at 4◻°C whereas all other antibodies were incubated for 1h at room temperature. Muscle bundles were stained for titin as previously described ^60^. Imaging of both 2D and 3D muscle models were carried out using the CellInsight CX7 microscope and a cell-based analysis was carried out using the accompanied HCS Platform software (Thermo Fisher Scientific). 3D muscle bundles were imaged with 20X objective using confocal for 40 stacks at 5 μm, images show maximum projections. Fusion index indicates the percentage of segmented nuclei within MyHC positive object, the myotubes, compared to the total number of nuclei.

##### Calcium flux staining, imaging and quantification

Calcium flux analysis was carried out in immortalized muscle cells 96h after transfection, allowing mature myotubes to form but in contrast to 120h after transfection, stay fully attached. Fluo-4 calcium imaging kit (Invitrogen) was then used according to the manufacturer’s instructions. Neuro background suppressor was added 1:5 in 50 μL normal medium before imaging. Live cell imaging was carried out with the BZ-X800 Live-Cell Imaging Microscope (Keyence) using the 20X objective. The fluorescent signal was recorded for 30 seconds prior to KCl stimulation, which was done by directly adding 50 μL of normal medium containing 20 μM KCl to the well. Live cell imaging was stopped after circa 45 seconds of KCl stimulation. Analysis was performed using CALIMA ^62^. Average calcium fluxes were calculated using cell-based analysis using CALIMA software. Only adherent cells were imaged and quantified. Myoblasts or myotubes from 3 wells were segmented based on the MFI using the CALIMA software.

#### mRNA expression analyses

RNA was extracted using Qiazol, precipitated using chloroform followed by RNA isolation using the PureLink™ RNA Mini Kit (Invitrogen) according to the manual. cDNA synthesis was carried out with the RevertAid H Minus First Strand cDNA Synthesis Kit according to the user guide (Thermo Fisher Scientific). Quantitative RT-qPCR was performed using SYBR® Green Master mix (Bio-Rad). Samples were run in biological and technical triplicates using the LightCycler 480 System (Roche Diagnostics). The following primers were used to assess gene expression: *MEF2A* Fw ‘TGAAAGCAGAACCAACTCGGA’ and Rv ‘GTAGGACAAAGCATTGGGGC’; *MYH3* Fw ‘TCAAAGAGTTGCAGGCTCGAA’ and Rv ‘CATAGTCGCTGCGCTGTTTC’; *USP18* Fw ‘CAACGTGCCCTTGTTTGTC’ and Rv ‘TGCAGTCTCTCCACCAAGTG’;

*MYH8* Fw ‘ACATTACTGGCTGGCTGGAC’ and Rv ‘TTCGCGCTGCTATCTGCTTC’; *MYOD* Fw ‘CGACGGCATGATGGACTACA’ and Rv ‘GCAGTCTAGGCTCGACACC’; *MYOG* Fw ‘GCTCAGCTCCCTCAACCAG’ and Rv ‘GTGAGCAGATGATCCCCTGG’; and *TBP* as housekeeping gene with Fw ‘CGCCGAATATAATCCCAAGCG’ and Rv ‘CCTGTGCACACCATTTTCCC’.

##### RNA sequencing and analysis

Poly(A) library preparation and sequencing were carried out by GENEWIZ Germany GmbH as follows. RNA samples were quantified using Qubit 4.0 Fluorometer (Life Technologies, Carlsbad, CA, USA) and RNA integrity was checked with RNA Kit on Agilent 5300 Fragment Analyzer (Agilent Technologies, Palo Alto, CA, USA). RNA sequencing libraries were prepared using the NEBNext Ultra II RNA Library Prep Kit for Illumina following the manufacturer’s instructions (NEB, Ipswich, MA, USA). Briefly, mRNAs were first enriched with Oligo(dT) beads. Enriched mRNAs were fragmented for 15 minutes at 94 °C. First strand and second strand cDNAs were subsequently synthesized. cDNA fragments were end repaired and adenylated at 3’ends, and universal adapters were ligated to cDNA fragments, followed by index addition and library enrichment by limited-cycle PCR. Sequencing libraries were validated using NGS Kit on the Agilent 5300 Fragment Analyzer (Agilent Technologies, Palo Alto, CA, USA), and quantified by using Qubit 4.0 Fluorometer (Invitrogen, Carlsbad, CA). The sequencing libraries were multiplexed and loaded on the flowcell on the Illumina NovaSeq 6000 instrument according to manufacturer’s instructions. The samples were sequenced using a 2×150 Pair-End (PE) configuration v1.5. Image analysis and base calling were conducted by the NovaSeq Control Software v1.7 on the NovaSeq instrument. Raw sequence data (.bcl files) generated from Illumina NovaSeq was converted into fastq files and de-multiplexed using Illumina bcl2fastq program version 2.20. One mismatch was allowed for index sequence identification.

RNA-Seq files were processed using the opensource BIOWDL RNAseq pipeline v5.0.0 (https://zenodo.org/record/5109461#.Ya2yLFPMJhE) developed at the LUMC. This pipeline performs FASTQ preprocessing (including quality control, quality trimming, and adapter clipping), RNA-Seq alignment, read quantification, and optionally transcript assembly. FastQC was used for checking raw read QC. Adapter clipping was performed using Cutadapt (v2.10) with default settings. RNA-Seq reads’ alignment was performed using STAR (v2.7.5a) on the GRCh38 human reference genome. The gene read quantification was performed using HTSeq-count (v0.12.4) with setting “–stranded=no”. The gene annotation used for quantification was Ensembl version 104. Differentiation expression analysis was carried out with DESeq2 ^63^. P-values were corrected for multiple testing using the Benjamini-Hochberg procedure.

###### General statistics and bioinformatics

Statistical significance for experiments in cell cultures was assessed using a student’s t-test with N=3, unless depicted otherwise. Enrichment analyses were performed using DAVID bioinformatics resources v6.8 (https://david.ncifcrf.gov/) with the following settings: Medium confidence score, total proteome (N=3349) or total transcriptome (N=11631) as background. Subsequent protein-protein interaction networks were analysed using STRING (https://string-db.org/; v11.5) and visualised with Cytoscape (v.3.8.2). Only significantly enriched clusters (Bonferroni FDR<0.05) of N>5 were considered.

## Acknowledgements

We would like to thank the TDI MS Laboratory/Discovery Proteomics Facility for their technical support. We thank Anneke J. van der Kooi and Joost Raaphorst, Amsterdam University Medical Centre for discussions on possible USP18 role in myositis.

## Author contributions

C.S.O., V.R. and B.M.K. directed this study. C.S.O. conducted most experiments and analyses. A.P.F. helped with mass spectrometry sample preparation and data interpretation. A.D. helped with a cell culture experiment. H.M. analysed the RNAseq. B.H. performed gene expression analysis for muscle bundles. E.W. and J.G. helped with muscle bundle culture and contractile analysis. I.V. performed mass spectrometry. Manuscript was written by C.S.O, B.M.K and V.R.; All authors commented on the manuscript.

## Data availability

The mass spectrometry data set has been deposited to the ProteomeXchange Consortium (http://proteomecentral.proteomexchange.org) via the PRIDE partner repository ^64^ and is available through the dataset identifier PXD032406. The transcriptomics dataset is deposited in NCBI’s Gene Expression Omnibus (GEO) ^65^ and is accessible under the accession number GSE198794, https://www.ncbi.nlm.nih.gov/geo/query/acc.cgi?acc=GSE198794.

