## Supplementary material for "USP18 is an essential regulator of muscle cell differentiation and maturation"

This PDF file includes:

Supplementary Text

Figs. S1 to S8

Tables S1 to S4

### Fig. S1

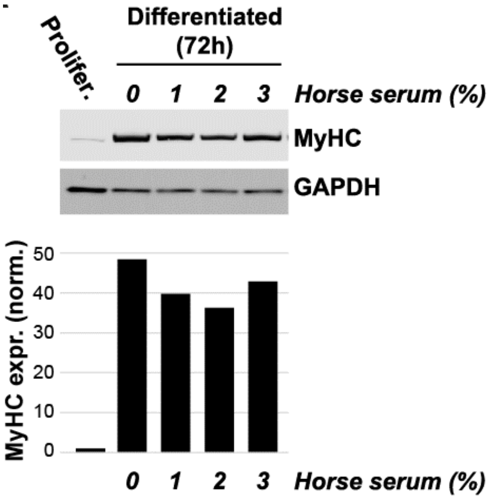


**Figure S1. Differentiation in presence or absence of horse serum rich medium.** Representative western blot shows MyHC expression after 72 hours of starvation induced differentiation with different horse sera content. GAPDH was used as loading control. Bar chart shows relative MyHC expression normalised to GAPDH.

### Fig. S2

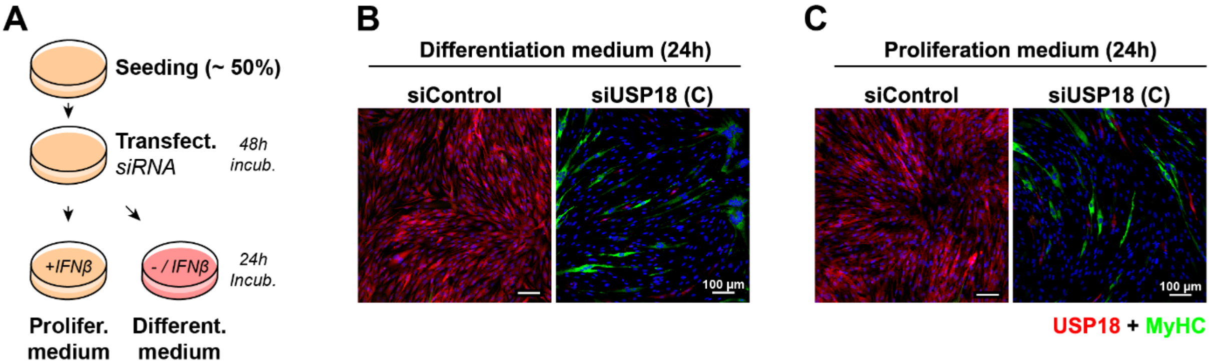


**Figure S2. USP18 siRNAs reduce USP18 expression** (**A**) Schematic representation of experimental setup. (**B-C**) Representative immunofluorescence images show MyHC (green) and USP18 (red) staining of cell transfected with siControl or siUSP18 cultured for 24h in differentiation (B) or proliferation (C) medium containing IFNβ (1000 U/mL). Nuclei are coloured in blue. Scale bar is 100 µm.

Fig. S3

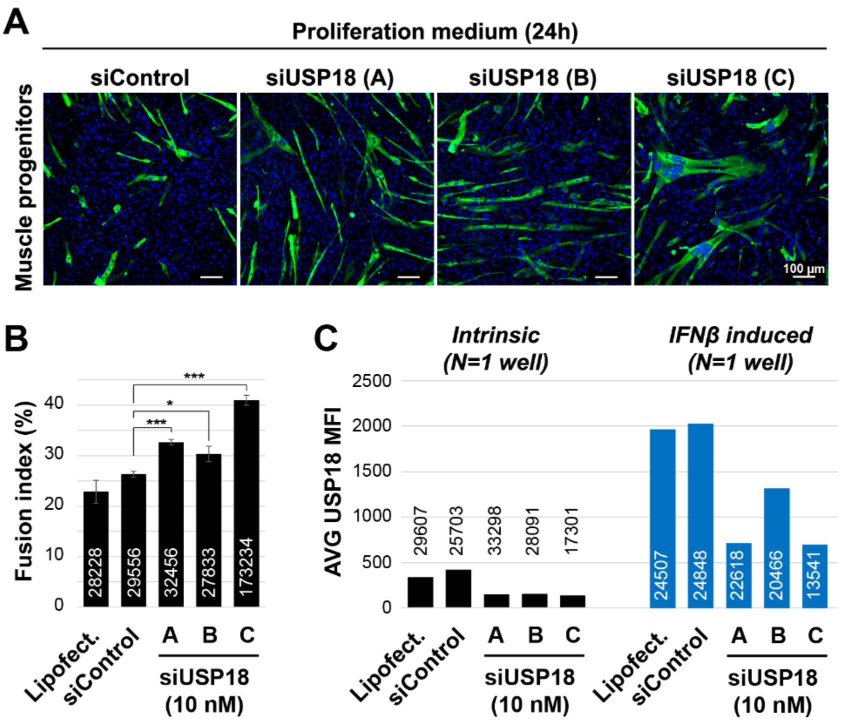


**Figure S3. USP18 siRNAs reduce USP18 expression and induce premature differentiation under proliferating conditions in muscle progenitors.** (**A**) Representative immunofluorescence images show MyHC staining in myogenic progenitor cells after 24 hour cultured in proliferation medium of three USP18 siRNAs. Nuclei are counterstained with DAPI (blue). Scale bar is 100 µm. (**B-C**) Bar charts show the fusion index (B) and the respective USP18 mean fluorescence intensity (MFI) (C) of muscle progenitors. USP18 MFI was measured from untreated or IFNβ-treated cell cultures (C). Number of cells is above/inside the bars. In panel C, average and error bars (SD) are from N=3. Statistical significance is depicted by asterisks. (*: 0.05-0.005, or *** <0.0005). Panel C was average MFI over the indicated number of cells of one well.

### Fig. S4

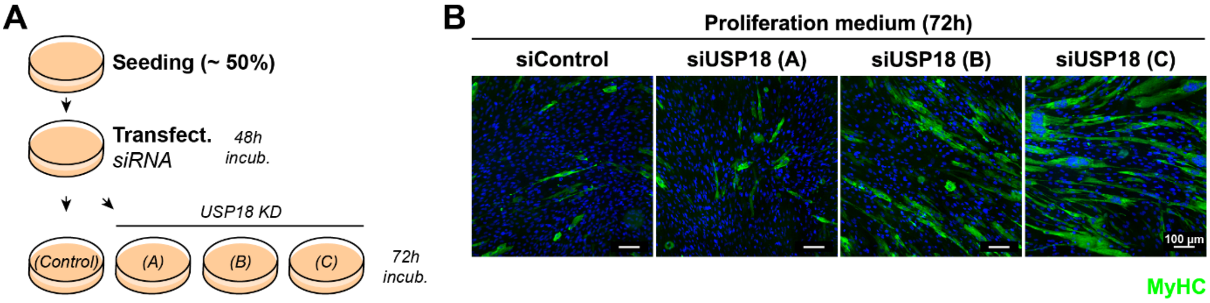


**Figure S4. Full premature differentiated muscle cells under proliferating conditions in immortalized muscle cells.** (**A**) Schematic representation of experimental setup. (**B**) Representative immunofluorescence images of siControl or siUSP18 transfected cultures 72 hours after incubation in proliferation medium show MyHC staining. Nuclei are counterstained with DAPI (blue). Scale bar is 100 µm.

### Fig. S5

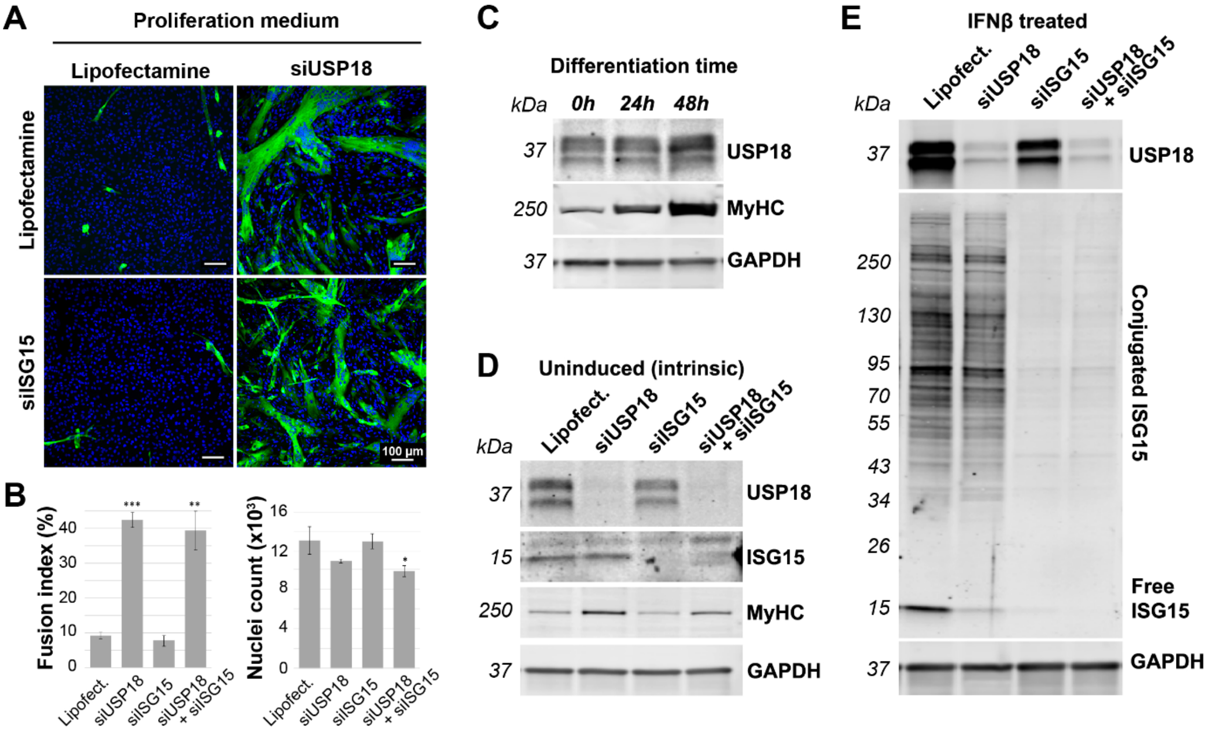


**Figure S5. USP18 KD induces differentiation reprograming in myogenic progenitors.** (**A**) Representative immunofluorescence images show differentiating cells in siUSP18 cultures and co-transfection with siISG15 in proliferation medium. Scale bar is 100 µm. (**B**) Bar charts show fusion index and the respective cell count. Average and error bars (SD) are from N=3 biological replicates. Statistical significance is depicted by asterisks. (*: 0.05-0.005, **: 0.005-0.0005, *** <0.0005). (**C**) Representative western blot shows USP18 and MyHC expression during differentiation. GAPDH indicates equal loading. (**D-E**) Representative western blots show USP18 and ISG15 expression. MyHC marks differentiation and GAPDH is a loading control. Panel B shows intrinsic (uninduced) protein expression and panel C shows expression after IFN-1 induction.

### Fig. S6

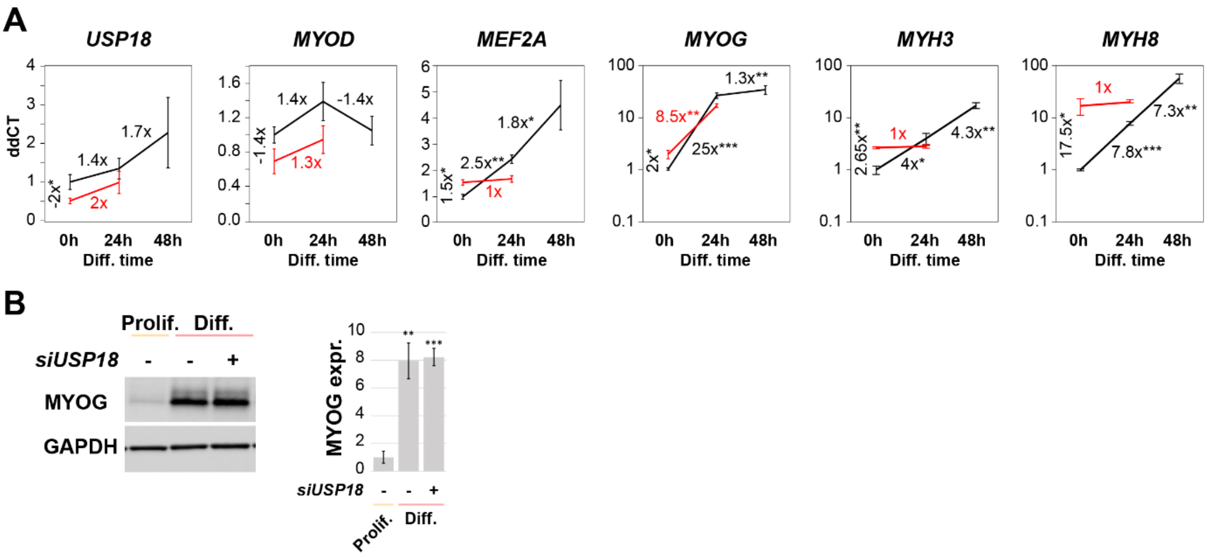


**Figure S6. USP18 KD re-programs the myogenic regulatory factors profile, RT-qPCR validation. (A)** Graphs show the average ddCT values for differentiation in control (black) and USP18 KD (red) cell cultures and the respective fold changes. **(B)** Representative western blot shows the MYOG expression. Bar chart shows the quantifications for which GAPDH was used as loading control. Expression levels were normalised to control samples in proliferation medium (Prolif.). Average and error bars (SD) are from N=3. Statistical significance is depicted by asterisks. (*: 0.05-0.005, **: 0.005-0.0005, *** <0.0005). Results were obtained in immortalized cell cultures.

**Fig. S7**

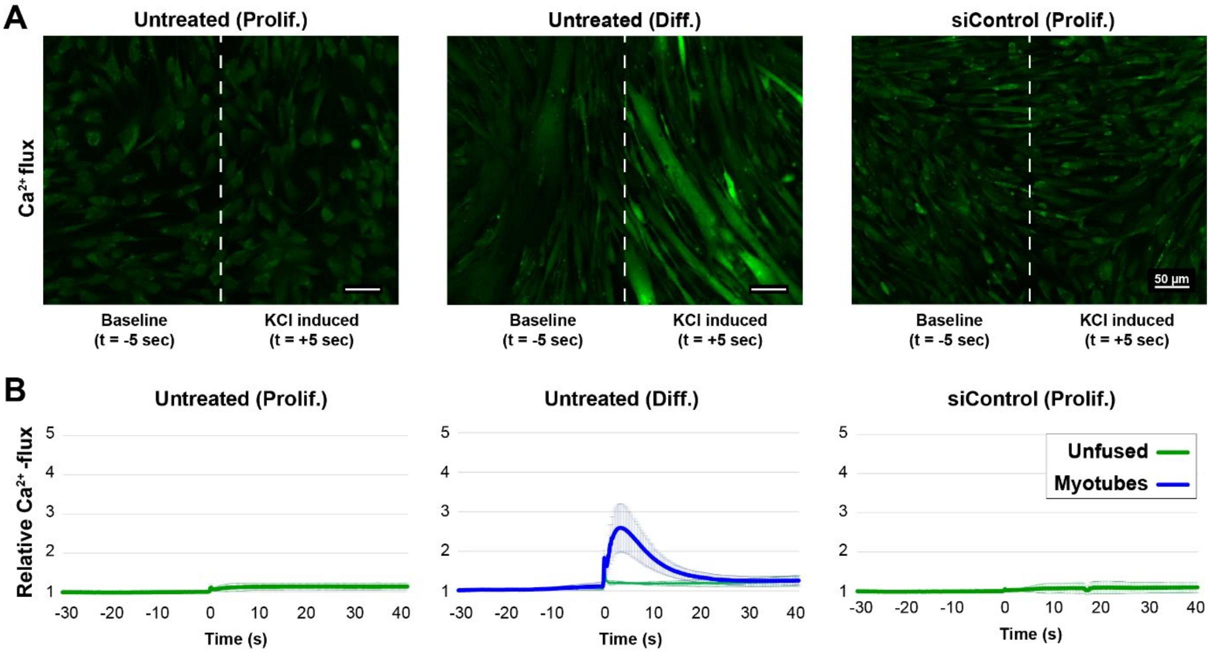


**Figure S7. Calcium flux for proliferating and differentiating immortalized muscle cells.** (**A**) Representative images show the calcium flux (green signal) 5 seconds before or after KCl (10 mM) administration in untreated or siControl transfected cells cultured in proliferation or differentiation medium. The respective images are separated by a white dashed line. Scale bar is 50 µm. (**B**) Graphs show the change in calcium flux over time. Calcium flux in uni-nucleated (unfused) or multi-nucleated (myotubes) cells is depicted by green or blue lines. Average and error bars (SD) are from N=3 biological replicates.

### Fig. S8

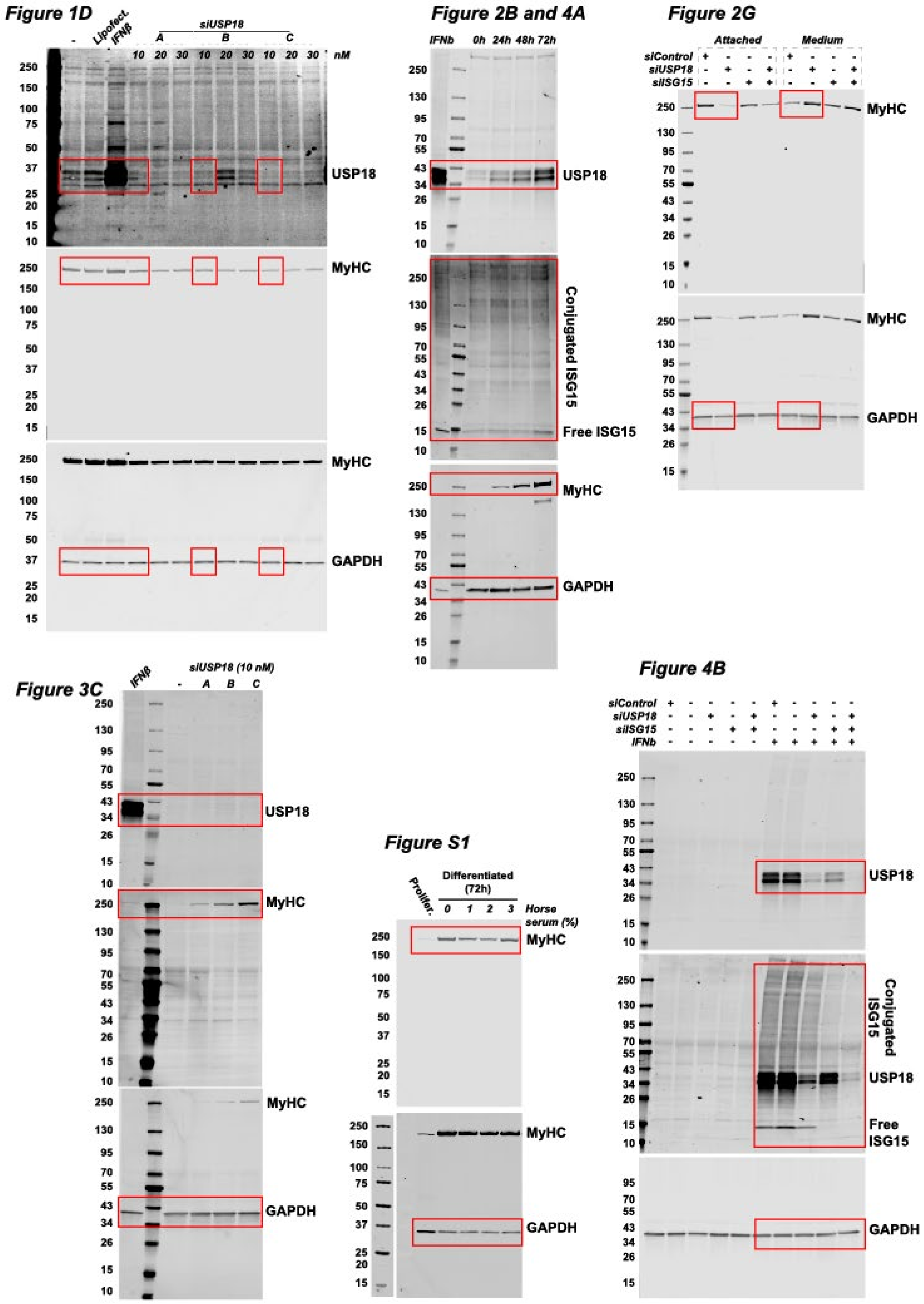
Figure 1D
Lipofect.
IFNβ
siUSP18
A B C
10 20 30 10 20 30 10 20 30 nM
USP18
MyHC
MyHC
GAPDH
Figure 2B and 4A
IFNb 0h 24h 48h 72h
USP18
Conjugated ISG15
Free ISG15
MyHC
GAPDH
Figure 2G
Attached
Medium
siControl
siUSP18
siISG15
MyHC
MyHC
GAPDH
Figure 3C
IFNβ
siUSP18 (10 nM)
A B C
USP18
MyHC
MyHC
GAPDH
Figure S1
Prolifer.
Differentiated (72h)
0 1 2 3
Horse serum (%)
MyHC
MyHC
GAPDH
Figure 4B
siControl
siUSP18
siISG15
IFNb
USP18
Conjugated ISG15
USP18
Free ISG15
GAPDH

### Figure 6

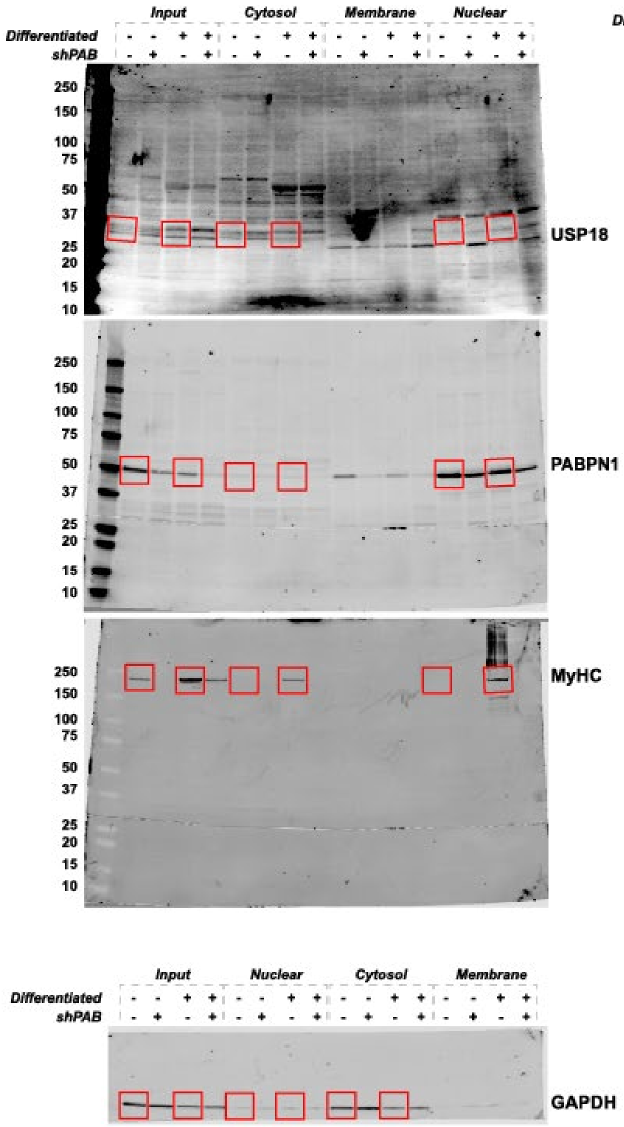


### Figure 7B and S6 (Myog)

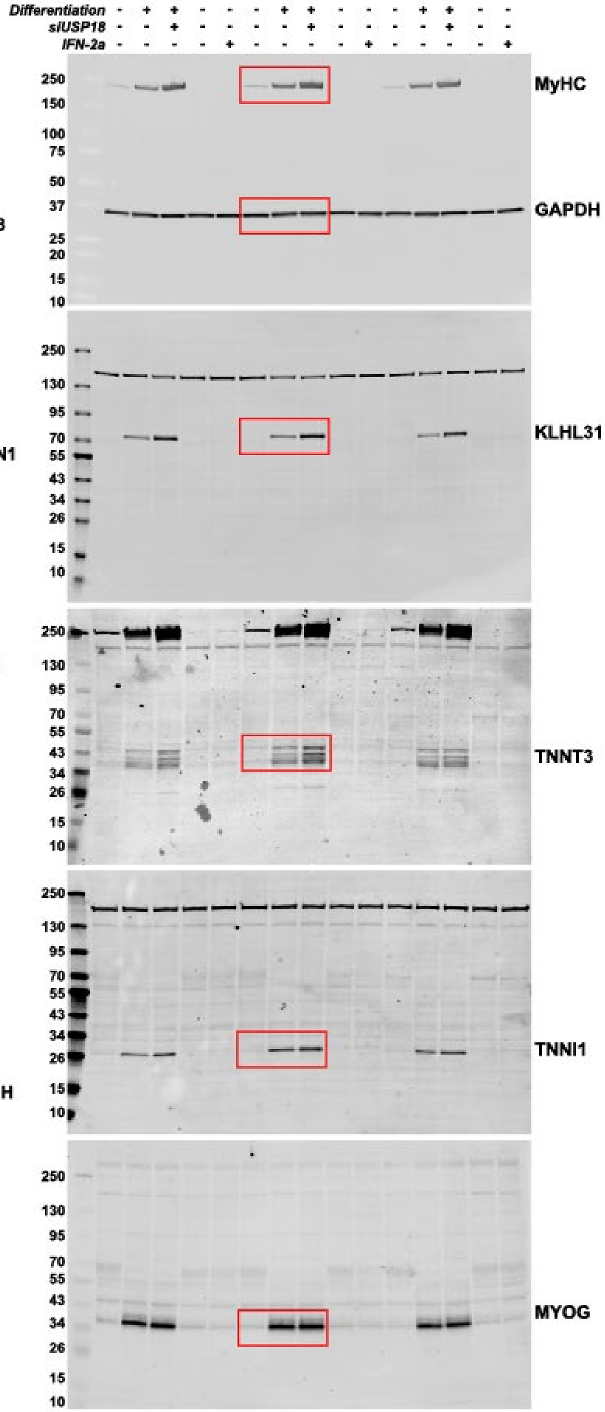

*Figure S5. miPSC*

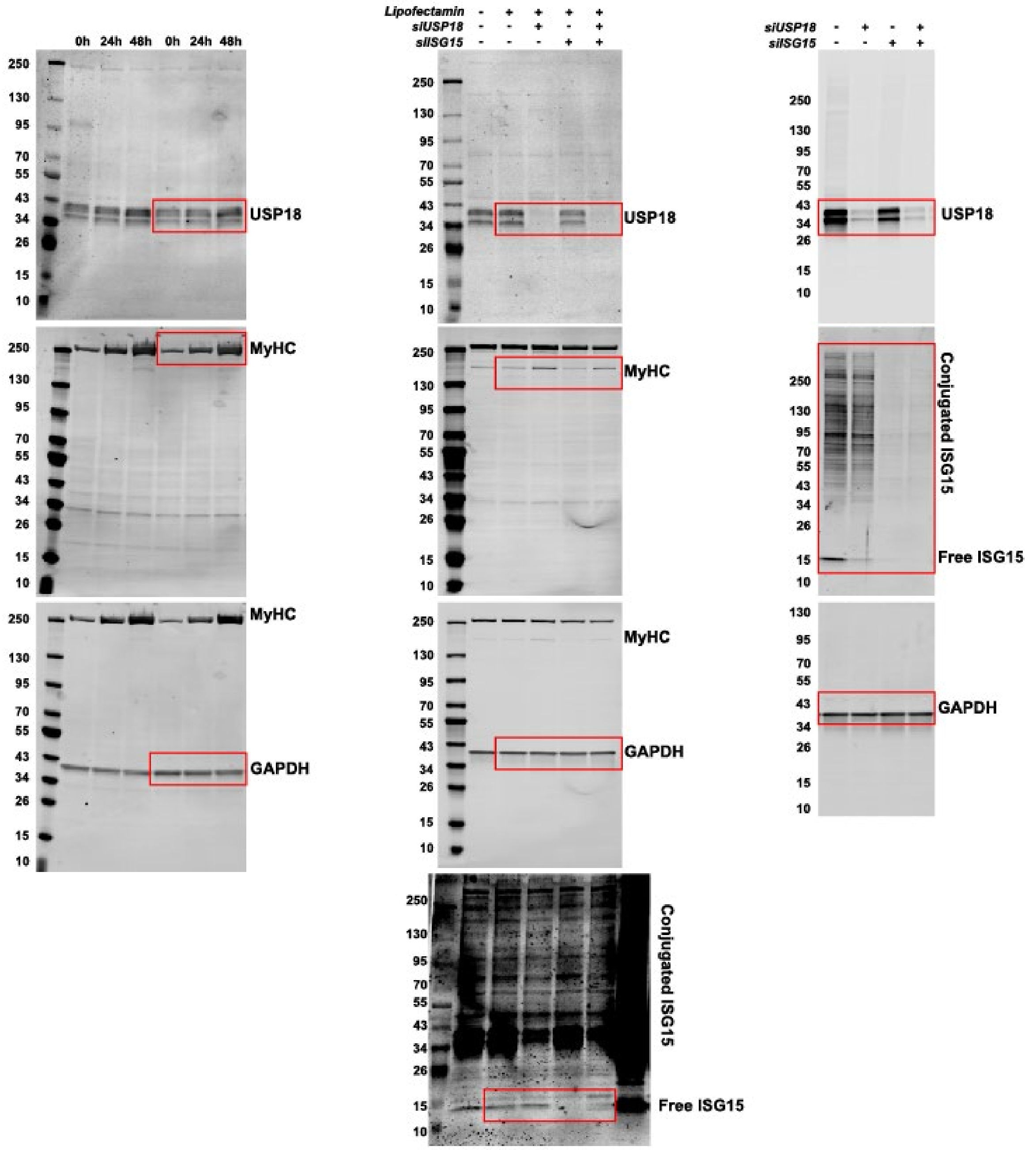


**Figure S8. Full uncropped western blots.** All western blots of the respective main and supplementary figures. Red boxes indicate the cropped areas of the blot as shown in those figures. Western blots are presented in order of antibody incubation, for instance first incubated with antibody against USP18, imaged in the 700nm channel and reblotted with antibody against MyHC and subsequently GAPDH in the 800nm channel. The blots of figures 1D, 2G, 3C, S1, 4B, 6 and some of figure 7B and S5 were performed in such a sequential order. For the remaining blots, protein lysate was blotted on multiple western blots and individually immunolabelled.

**Table S1.**

| siRNA | Supplier | Cat. number / sequence |
|---|---|---|
| siControl | Dharmacon | #D-001206-13 |
| USP18 (A) | Eurofins | CUGCAUAUCUUCUGGUUUA |
| USP18 (B) | Eurofins | GGAAGAAGACAGCAACAUG |
| USP18 (C) | Thermo Fisher | AM16708, ID: 105194 |
| ISG15 | Dharmacon | #L-004235-03-0005 |

**Table S1.** List of used siRNAs

**Table S2.**

| | Protein | Dilution | | Host | Supplier | Cat. number |
|---|---|---|---|---|---|---|
| | | IF | WB | | | |
| Primary antibodies | USP18 | 1:100 | 1:1000 | anti-Rabbit | Cell signaling | #4813 |
| | ISG15 | 1:100 | 1:1000 | anti-Rabbit | Cell signaling | #2743 |
| | pSTAT1 | - | 1:1000 | anti-Rabbit | Cell signaling | #9167 |
| | GAPDH | - | 1:5000 | anti-Mouse | Invitrogen | MA5-15738 |
| | KLHL31 | - | 1:1000 | anti-Rabbit | ProteinTech | 24781-1-AP |
| | TNNI1 | - | 1:1000 | anti-Rabbit | Novus Biologicals | NBP1-56641 |
| | MyHC | 1:250 | 1:250 | anti-Mouse | DSHB | MF20 |
| | TNNT3 | - | 1:1000 | anti-Mouse | DSHB | JLT12 |
| | TTN | 1:50 | - | anti-Mouse | DSHB | 9D10 |

| | Target | Fluorophore | Dil. | Supplier |
|---|---|---|---|---|
| 2nd antibodies | Anti-Rabbit | Cy5 | 1:5000 | Thermo Fisher Scientific |
| | Anti-Mouse (IgG or IgM) | 488 | 1:5000 | Thermo Fisher Scientific |
| | WB anti-Rb/M | 800/680 | 1:10000 | IRDye 800CW or IRDye 680RD (LI-COR) |

**Table S2.** List of used antibodies for immunofluorescence and western blotting.

**Table S3.** The USP18-dependent Transcriptome. Related to Fig. 3F and Fig. 5-6

**Table S4.** The USP18-dependent Proteome. Related to Fig. 7
